# Astrocyte-neuron interplay is critical for Alzheimer’s disease pathogenesis and is rescued by TRPA1 channel blockade

**DOI:** 10.1101/2021.03.29.437466

**Authors:** Adrien Paumier, Sylvie Boisseau, Karin Pernet-Gallay, Alain Buisson, Mireille Albrieux

## Abstract

**Background:** The sequence of cellular dysfunctions in preclinical Alzheimer’s disease must be understood if we are to plot new therapeutic routes. Hippocampal neuronal hyperactivity is one of the earliest events occurring during the preclinical stages of Alzheimer’s disease in both humans and mouse models. The most common hypothesis describes amyloid β accumulation as the triggering factor of the disease but the effects of such accumulation and the cascade of events leading to cognitive decline remain unclear. In mice, we previously showed that amyloid β-dependent TRPA1 channel activation triggers hippocampal astrocyte hyperactivity, subsequently inducing hyperactivity in nearby neurons. In this work, we investigated the potential protection brought by an early chronic pharmacological inhibition of TRPA1 channel on Alzheimer’s disease progression.

**Methods:** We administered a specific inhibitor of TRPA1 channel (HC030031) intraperitoneally from the onset of amyloid β overproduction in the APP/PS1-21 mouse model of Alzheimer’s disease. We characterized short-, medium-, and long-term effects of this chronic pharmacological TRPA1 blockade on Alzheimer’s disease progression at functional (astrocytic and neuronal activity), structural, biochemical, and behavioural levels.

**Results:** Our results revealed that the first observable disruptions in the Alzheimer’s disease transgenic mouse model used correspond to aberrant hippocampal astrocyte and neuron hyperactivity. We showed that chronic TRPA1 blockade normalizes astrocytic activity, avoids perisynaptic astrocytic process withdrawal, prevents neuronal dysfunction, preserves structural synaptic integrity and strengthens the glial plaque barrier. These protective effects preserved spatial working-memory in this Alzheimer’s disease mouse model.

**Conclusion:** The toxic effect of amyloid β on astrocytes triggered by TRPA1 channel activation is pivotal to Alzheimer’s disease progression. TRPA1 blockade prevent irreversible neuronal dysfunction, making this channel a potential therapeutic target to promote neuroprotection.

## Introduction

Alzheimer’s disease starts long before its clinical diagnosis. The primary mechanisms leading to the onset of the disease must be understood if we are to achieve successful medical control, delaying or even reversing the transition from normal brain physiology to cognitive impairments. More than two decades ago, amyloid-β (Aβ) dyshomeostasis was proposed to be the major initiating factor in Alzheimer’s disease, occurring upstream of alterations in other proteins^1^. Aβ amyloidosis was associated with functional changes in the brain, such as impaired neuronal activity^2^. Several lines of evidence indicate that neuronal hyperactivity may be a key feature in early-stage Alzheimer’s disease^3, 4^. Thus, functional human imaging studies^5^ revealed the prodromal phase of Alzheimer’s disease to be characterized by neuronal hyperactivity in the hippocampus, associated with amnestic mild cognitive impairment (aMCI), where memory deficits are present but do not significantly impair routine activity^5^. In Alzheimer’s disease mouse models, early excessive neuronal activation within the hippocampus was shown to be linked to the presence of soluble oligomeric Aβ^3^, and to crucially contribute to cognitive decline and disease progression^4, 6, 7^.

In the brain, neuronal activity is tightly regulated by astrocytes as they sense the activity taking place in the synapses they enwrap. In these tripartite synapses, astrocytes can modulate neurotransmission at the synapse by taking up or releasing neuroactive factors^8^. These key cells react to virtually all types of pathological alterations and have been implicated in the pathogenesis of several neurodegenerative disease, including Alzheimer’s disease ^9^, where they have been reported to become hyperactive^10–12^. In the early stages of Alzheimer’s disease in murine models, our group^12^ showed that this astrocytic hyperactivity is present from the onset of Aβ production, well before inflammation becomes discernible, and that it drives the initial neuronal hyperactivity within the CA1 hippocampus. We identified an astrocytic calcium channel (TRPA1, transient receptor potential ankyrin 1) implicated in this early toxicity and showed that specific acute pharmacological blockade of TRPA1 reversed the abnormal activities of both astrocytes and neurons^12^.

TRPA1 is a nonselective calcium channel that was initially described in nociceptive peripheral sensory neurons^13^. Within the central nervous system, TRPA1 is expressed on astrocytes in the hippocampus, is involved to some extent in basal Ca^2+^ signalling in physiological conditions, and seems to behave as an “aggression sensor” in noxious conditions^14–17^. A specific TRPA1 inhibitor (HC030031) can be easily delivered intraperitoneally, and has been successfully used to treat neuropathic pain^18–20^. This specific inhibitor traverses the blood brain barrier and no adverse effects were reported in human preclinical studies^21^.

With this study, we aimed to assess the potential protection afforded by chronic pharmacological inhibition of TRPA1 and to characterize its short-, medium-, and long-term effects on Alzheimer’s disease progression at the functional, morphological, biochemical, and behavioural levels. We found that chronic treatment with TRPA1 inhibitor has a strong beneficial effect on Alzheimer’s disease progression at multiple levels. It normalized astrocytic and neuronal activity while protecting structural synaptic integrity from irreversible degeneration, improving astroglial plaque barrier function and partly preventing mnesic decline.

## Materials and methods

### Animals

All experiments were performed according to animal care guidelines developed based on the European Community Council directives of November 24, 1986 (86/609/EEC), in accordance with French national institutional animal care guidelines (protocol APAFIS#19142-2018121715504298). The study protocol was approved by the “Grenoble Institute of Neurosciences Ethics Committee” and followed the ARRIVE guidelines. The transgenic mouse model used co-expresses the KM670/671NL mutated human form of amyloid precursor protein (mutated APP) and the L166P form of presenilin 1 (mutated PS1) under the control of the Thy1 promoter^22^. This mouse line was maintained in a heterozygous state and backcrossed onto a C57Bl6/J genetic background (Janvier Labs, France). In some experiments, APP/PS1-21 mice were cross-bred with the transgenic Thy1-eYFP-H^23^ mouse, expressing fluorescent protein (eYFP) in specific populations of projection neurons. Animals were housed in groups on a 12 h light/dark cycle with food and water available *ad libitum*.

### Chronic intra-peritoneal drug treatment

The TRPA1-specific inhibitor HC030031^18^ (Tocris Bioscience) was solubilized in 16% DMSO, 0.6% Pluronic acid F-127, and sterile saline. HC030031 (IC_50_ ≈ 7.6 µM^18^) was administered intraperitoneally every day (5 mg/kg body weight) to male and female transgenic mice and their wild-type age-matched and sex-matched littermates (WT) from the 14th postnatal day. At this time, Thy1 gene promotor expression starts in the hippocampus^23^, triggering overproduction of Aβ in the APP/PS1-21 model. The same dose was previously used in models of neuropathic pain in rodents, and induced a substantial analgesic effect without toxicity^18–20^. The effects of HC030031 treatment were compared to those of vehicle treatment, administered according to the same schedule, in matched groups of APP/PS1-21 and WT groups. As births occurred, animals from each litter were included in the different groups randomly, balancing the sexes and the genotypes.

### Slice preparation

Coronal brain or sagittal hippocampal slices (300 µm thick) were prepared from APP/PS1-21 transgenic mice (1- and 3-month-old) or age-matched and sex-matched WT littermates. Mice were killed by decerebration and decapitated. The brain was rapidly removed for cutting in ice-cold cutting artificial cerebrospinal fluid (ACSF) made up of: 2.5 mM KCl, 7 mM MgCl_2_, 0.5 mM CaCl_2_, 1.2 mM NaH_2_PO_4_, 25 mM NaHCO_3_, 11 mM D-glucose, and 234 mM sucrose then cut using a vibratome VT1200S (Leica, Wetzlar, Germany). The solution was continuously bubbled with 95% O_2_ and 5% CO_2_. Slices containing the hippocampus were placed in ACSF composed of: 126 mM NaCl, 2.5 mM KCl, 1.2 mM MgCl_2_, 2.5 mM CaCl_2_, 1.2 mM NaH_2_PO_4_. Solutions were supplemented with 1 mM sodium pyruvate and continuously bubbled with 95% O_2_ and 5% CO_2_ at room temperature for a recovery period. Where indicated, slices were bathed in 40 µM HC030031 for 5 min before commencing and throughout calcium imaging or electrophysiological recordings.

### Single-astrocyte dye-loading with Fluo-4

Coronal brain slices were transferred to a chamber at room temperature allowing constant perfusion with ACSF bubbled with 95% O_2_ and 5% CO_2_ on the stage of an upright compound microscope (Eclipse E600 FN, Nikon, Paris, France). The microscope was equipped with a water-immersion 60x objective (NA 1.0) and infrared differential interference contrast optics with CCD camera (Optronis VX45, Kehl, Germany). Glass pipettes 8-11 MΩ (Harvard apparatus) were filled with intracellular solution consisting of: 105 mM K-gluconate, 30 mM KCl, 10 mM phosphocreatine, 10 mM HEPES, 4 mM ATP-Mg, 0.3 mM GTP-Tris, 0.2 mM Fluo-4 pentapotassium salt (Life Technologies), adjusted to pH 7.2 with KOH. Signals were amplified by Axopatch 200B, sampled by a Digidata 1440A interface and recorded with pClamp10 software (Molecular Devices, Foster City, USA). Astrocytes were identified based on their morphology, localization in the *stratum radiatum* and negative resting potential (between - 70 and -80 mV). The membrane potential was held at -80 mV. Input resistance was calculated based on the current measured following a 10-mV pulse lasting 80 ms, near the end of the voltage command. Only passive astrocytes showing a linear *I/V* relationship and low input resistance (∼ 50 MΩ) were retained for dye loading. Once the whole-cell configuration had been stabilized, access resistance was constantly monitored. Astrocytes were excluded from further study when this parameter varied > 20% during the experiment. To allow sufficient diffusion of the dye and avoid astrocyte dialysis, the time in whole-cell configuration was limited to less than 5 min. After this period, the patch pipette was carefully withdrawn to allow the astrocyte to recover. To maximize diffusion of the dye into the astrocytic processes, a lapse of at least 5 min was allowed before starting calcium imaging^12, 24, 25^.

### Calcium imaging

Single-astrocyte dye-loaded slices were placed in a constantly perfused chamber on the stage of an upright compound microscope (Eclipse E600 FN, Nikon, Paris, France) equipped with a water-immersion 60x (NA 1.0) objective and a confocal head (confocal C1 head, Nikon, Paris, France). Fluorescent images were recorded following excitation at 488 nm; emission was detected with a 515 ± 15 nm filter. Images were acquired with EZ-C1 software (Nikon, Paris, France) at 1.2-s intervals in a single confocal plane over a 5-min period.

Ca^2+^ transients were measured in two-dimensional images, in individual subregions matching the shape of the astrocyte structure. Manually selected ROIs (∼1 µm²) were placed along astrocytic processes lying in the focal plane^12, 24^ and a ROI was also selected in the soma if accessible. Prior to analysis, raw images were stabilized (if slight *x-y* drift occurred during recordings, z drifts were excluded) using the ImageJ plugins Template Matching and Filtered, and applying a 3D Hybrid Median Filter^26^. Intracellular Ca^2+^ activity was measured by analyzing the fluorescence signal F within each ROI, using *CalSignal* software^27^. Significant changes in fluorescence were detected on the basis of the ΔF/F_0_ ratios. F_0_ was determined for each ROI throughout the recording period, and a Ca^2+^ event was defined as a significant and continuous signal increase exceeding a fixed threshold followed by a significant and continuous signal decrease exceeding the same threshold. Thus, ROIs were defined as active when fluorescence increased ≥ 2 standard deviations relative to baseline. After automated peak detection, each Ca^2+^ transient was visually checked by an operator.

### Electrophysiological recordings

Whole-cell recordings were made in the somata of visually identified CA1 pyramidal neurons. Patch pipettes (4-6 MΩ) were filled with an internal solution composed of: 105 mM K-gluconate, 30 mM KCl, 10 mM phosphocreatine, 10 mM HEPES, 4 mM ATP-Mg, 0.3 mM GTP-Tris, 0.3 mM EGTA, adjusted to pH 7.2 with KOH. Spontaneous excitatory post-synaptic currents (sEPSCs) were recorded at a membrane holding potential of -60 mV. All currents were recorded at room temperature (22-24°C), and only one neuron was studied in each slice. sEPSCs and their kinetics were analysed within 5 min of recording, after a stabilization period of 10 min. Access resistance was constantly monitored, and recordings were excluded from further study if this value varied by more than 20% during the experiment. Recordings were analysed using the Clampfit module in pClamp8 software (Molecular Devices, Foster City, USA). A threshold of 20 pA was set to exclude miniature sEPSCs. Events were accepted as slow inward currents (SICs) if their time-to-peak exceeded 20 ms (rise time) and their amplitude was > 20 pA^28^.

Where indicated, Aβo was prepared from recombinant Aβ_1-42_ peptide (Bachem) and hippocampal slices were perfused with 100 nM of Aβo for 5 min before recording ^12^. sEPSCs/SICs, and their kinetics were analysed in 5-min epochs within the time-frame of the recordings. The recording for each epoch was compared to an initial 5-min recording and sEPSCs were normalized relative to this initial value.

In some experiments, recombinant *R. gracilis* D-amino acid oxidase (DAAO, kindly provided by Jean-Pierre Mothet, Paris, France) was applied to brain slices (0.2 U/ml in ACSF) for at least 45 min to degrade D-serine. Slices were then perfused with DAAO-containing ACSF (0.2 U/ml) throughout imaging^29^.

### BAPTA dialysis

For BAPTA dialysis, pipettes were filled with the following solution: 100 mM K-gluconate, 30 mM KCl, 10 mM phosphocreatine, 10 mM HEPES, 4 mM ATP-Mg, 0.3 mM GTP-Tris, 0.15% biocytin and 10 mM BAPTA. Astrocytes were first passively loaded with BAPTA and biocytin for 20 min in voltage-clamp mode^30, 31^. Successive whole-cell recordings were performed in CA1 neurons as described above. Slices containing biocytin-filled cells were transferred to a freshly-prepared solution of 4% paraformaldehyde in 0.1 M phosphate buffered saline (PBS) and fixed overnight at 4°C. Biocytin was detected by incubating brain slices in Alexa 488-conjugated streptavidin (Life Technology, USA; 1:1000)^32^. Sections were washed with PBS, mounted in Dako fluorescent mounting medium (Dako, USA), then examined under an upright confocal microscope (C1, Nikon, France) to estimate dye diffusion.

### NMDAR D-serine site occupancy

Field excitatory post-synaptic potentials (fEPSPs) were recorded in CA1 *stratum radiatum* from hippocampal slices with a standard patch pipette filled with an extracellular solution containing 0.2 mM MgCl_2_ and supplemented with 10 µM 2,3-dihydroxy-6-nitro-7-sulphamoyl-benzo quinoxaline (NBQX) to isolate NMDAR fEPSPs. Synaptic responses were evoked by orthodromic stimulation (100 µs, 50-100 µA) of Schaffer collaterals induced using a bipolar stimulation electrode (Frederick Haer & Co., USA). The amplitude of NMDAR fEPSPs was monitored for 10 min to determine the basal response for each slice. As indicated, in some experiments, slices were perfused with 100 nM Aβo for 5 min before resuming recording for 10 min. NMDAR D-serine site occupancy was finally evaluated by adding 10 µM D-serine to determine maximal NMDAR fEPSPs potentiation^33^.

### Dendritic spine analysis

Hippocampal sections from Thy1-eYFP-H-APP/PS1-21 mice were imaged using a Zeiss Airyscan module with an oil-immersion Plan Apochromat 100x objective (NA 1.4). Confocal image stacks (200-nm step) were acquired with a voxel size of 0.041 × 0.041 × 0.2 µm. 3D analysis of spine density, spine volume and spine classification were performed using NeuronStudio software (Icahn School of Medicine at Mount Sinai, New York, USA). The length of individual dendrites was automatically measured, as was the number and volume of associated spines. Spines were classified as stubby, thin, or mushroom populations using automated 3D shape classification^34^. For each condition, 10 dendrites were imaged and 700 spines were analysed.

### Electron microscopy

Hippocampi were dissected before fixing with 1.7% glutaraldehyde in 0.1 M PBS pH 7.4 for 48 h at room temperature. The CA1 area was dissected under a binocular microscope and further fixed for 2 h in the same solution. Samples were then washed with buffer and post-fixed with 1% osmium tetroxide and 0.1 M PBS pH 7.2 for 1 h at 4°C. After extensive rinsing with water, cells were stained with 1% uranyl acetate (pH 4) in water for 1 h at 4°C before dehydration in graded ethanol baths (30% - 60% - 90% - 100% -100% - 100%) and infiltration with a mix of 1/1 epon/ethanol 100% for 1 h, followed by several baths of fresh epoxy resin (Sigma-Aldrich, France) over 3 h. Finally, samples were included in a resin-filled capsule and allowed to polymerize for 72 h at 60 °C. Ultrathin (60 nm) sample sections were cut with an ultramicrotome (Leica, USA). Sections were post-stained with 5% uranyl acetate and 0.4% lead citrate. Images were acquired under a transmission electron microscope at 80 kV (JEOL 1200EX, Japan) using a digital camera (Veleta, SIS, Olympus, Germany) at 50 000x magnification. The presence of astrocytic processes in contact with an excitatory synapse was estimated in the neuropil of the *stratum radiatum* from 42 randomly-selected areas in four sections from each of three mice for each condition. Astrocyte processes were identified by their relatively clear cytoplasm, angular shape compared to their smoother neuronal counterparts, and based on the presence of glycogen granules^35^.

### Immunohistochemistry

Deep anaesthesia was induced by intra-peritoneal injection of 320 mg/kg sodium pentobarbital. Mice were then perfused intracardially with 25 mL 0.1 M PBS followed by 25 mL 4% paraformaldehyde in 0.1 M PBS, pH 7.3. Brains were rapidly removed, post-fixed overnight at 4°C in 4% paraformaldehyde, immersed in 20% sucrose in 0.1 M PBS, pH 7.5 overnight, frozen in cooled (-35°C) isopentane, and stored at -30 °C. Serial frontal sections (20 µm thick) were cut with a cryostat microtome (HM 500 M, Microm, Francheville, France). Sections were blocked by incubation for 30 min in 3% bovine serum albumin in Tris-Buffered Saline (TBS)-Tween-Triton (TBSTT) (0.1 M Tris Base, 0.15 M NaCl, 0.1% Tween, 0.1% Triton X-100) (dilution/blocking buffer). Tissue sections were then incubated overnight at 4 °C with one of the following antibodies: anti-amyloid fibrils antibody^36^ (OC, Rockland, USA, rabbit polyclonal; 1:500), anti-GFAP antibody (Sigma, France, mouse monoclonal; 1:500) or anti-Iba1 antibody (Novus, USA, goat polyclonal; 1:500). Tissue sections were washed in TBSTT and incubated for 2 h at room temperature with Alexa 594-(Life Technology, USA; 1:1000), Alexa 488-(Life Technology, USA; 1:1000), or Alexa 647-(Life Technology, USA; 1:1000) conjugated secondary antibodies. Sections were washed in TBSTT, incubated for 5 min with Hoechst (33258, Sigma) to stain nuclei, and mounted in Dako fluorescent mounting medium (Dako, USA).

Where indicated, amyloid-β deposits were stained with thioflavin S^37^. Sections were re-hydrated in TBS buffer (0.1 M Tris Base, 0.15 M NaCl) and then incubated in filtered 1% aqueous thioflavin S (Sigma, France) for 8 min at room temperature in the dark, before washing several times in TBS.

### Image acquisition and data analysis

Sections were examined under a Zeiss LSM 710 confocal laser scanning microscope with an oil-immersion Plan Neofluar 40x objective (NA 1.3) or an oil-immersion Plan Neofluar 63x objective (NA 1.46) combined with a Zeiss Airyscan module to improve lateral resolution (∼140 nm) and signal-to-noise ratios. All images were recorded using the same exposure settings. For each immunostain, maximum intensity projections of 12 consecutive confocal *z* slices were processed in ImageJ. For quantitative analysis of thioflavin S and amyloid fibril (OC) staining, images were converted to 8-bit grey-level images, plaque area was determined by setting a fluorescence-intensity threshold (automated MaxEntropy algorithm) and measured using the Analyze and Measure functions in ImageJ^38^. The mean intensity for each plaque was calculated from the mean for all stained voxels. For quantitative analysis of GFAP and Iba1 processes stained around and within amyloid plaques, a ROI was defined to delimit the plaque. An 8-bit grey-level image was then created, to which a threshold was applied (automated Otsu algorithm) allowing the area of GFAP and Iba1 staining to be measured inside the ROI using the Analyze and Measure functions in ImageJ^39^.

### Dot-blot assays

Dissected hippocampi from 1-, 3- and 6-month-old APP/PS1-21 mice were homogenized in cold buffer containing 20 mM Tris-HCl (pH 7.5), 150 mM NaCl, 1 mM Na_2_EDTA, 1 mM EGTA, 1% NP-40, protease inhibitor cocktail (P8340, Sigma-Aldrich, France) and phosphatase inhibitors (P5726, Sigma-Aldrich, France). Samples were maintained at 4°C throughout experiments. Homogenates were cleared at 1000 x g for 10 min to remove nuclei and large debris. Loading buffer was added and samples were boiled for 10 min before resolving equivalent amounts of protein (20 µg, quantified by micro-BCA assay (Pierce) in duplicate extracts) on 4-20% gradient Bis-Tris polyacrylamide stain-free pre-cast gels (Bio-Rad) in denaturing conditions. Equivalent protein amounts (10 µg) were spotted onto nitrocellulose membranes using a Bio-Dot Microfiltration Apparatus (Bio-Rad). Proteins were first detected on membranes by Ponceau Red staining. Then, membranes were blocked with 3% bovine serum albumin (BSA) in TBST for 30 min at room temperature and incubated with anti-amyloid fibrils (OC) antibody (Rockland, USA, rabbit polyclonal, 1:1000) diluted in blocking solution, at 4 °C overnight. After several washes in TBST, blots were probed with HRP-conjugated anti-rabbit IgG (Fab’2) (Interchim, France; 1:40 000) antibody diluted in TBST for 45 min at room temperature. Finally, chemiluminescence was detected using the ECL detection system (Euromedex) and a ChemiDoc MP Imaging System (Bio-Rad). Chemiluminescence signals and total protein loading, based on Ponceau Red staining, were quantified using ImageJ software. Results are reported as the ratio of OC staining over total protein.

### Barnes maze experiments

The Barnes maze was initially designed to test spatial learning/memory in rodents; it takes advantage of rodents’ natural preferences for a dark environment^40^. Each mouse was assigned an escape box hole number that remained the same throughout the learning phase. The escape latency and the routes taken by the animal to reach the escape box were recorded to assess spatial learning. To evaluate reference memory, a probe test was performed by removing the escape box. Briefly, mice were placed in the centre of a brightly-lit open platform (BioSeb, diameter 120 cm). Visual spatial cues (4 large coloured shapes) were placed around the maze. The platform was confined with 20 holes (diameter 5 cm), with a hidden escape box (target) attached below one hole. On the first day (habituation), mice were placed in the middle of the platform and gently guided to the target after 5 min free exploration. Mice were left in the escape box for 1 min before being returned to their home cage. Over the three days of training/acquisition, mice performed three trials per day with an intertrial interval of 20 min. The mice had 180 s to find the escape box, after that time they were gently guided to it. On day 4 (probe test), the escape box was removed and mice were allowed to explore the maze freely for 60 s. The primary time to target and the number of explorations of the target area (target hole +/- 1) were analysed. All data were recorded and analysed with EthoVision XT9 (Noldus).

### Statistical analysis

The sample size for the various sets of data is mentioned in the corresponding figure legends. Results were expressed as mean ± SEM from independent biological samples; the distribution of experimental points is also shown. Data were analyzed using GraphPad Prism 9.0.0 software. Statistical comparisons between groups were conducted using the two-tailed Mann-Whitney test. For multiple comparisons, a Kruskal-Wallis test followed by Dunn’s multiple comparison test was used. Time-courses of cumulative frequency for SICs were analysed using the Kolmogorov-Smirnov test. The effects of time combined with treatment on Barnes maze learning were analysed using a 2-way ANOVA. Proportions of tripartite synapses were compared by applying Fischer’s exact test. Possible sex-dependence of neuronal activity, astrocytic activity, and mnesic performance was monitored in treated groups. No significant difference was found, therefore results for male and female animals were pooled for analysis (Supplementary Tables 1 and 2). Significance levels are indicated as follows in graphs: *, *p* < 0.05; **, *p* < 0.01; ***, *p* < 0.001 and n.s., not significant.

### Data availability

The authors confirm that the data supporting the findings of this study are available within the article and its supplementary material or are available from the corresponding author, upon reasonable request.

## Results

### Functional and structural alterations to neurons and astrocytes in the early and intermediate stages of Alzheimer’s disease

To decipher the early events in Aβ-induced pathophysiology, we first analysed the changes to neuronal and astrocytic activity in the hippocampus of APP/PS1-21 mice at 1 and 3 months old. These transgenic mice overexpress mutant APP in combination with mutant PS1, thus producing high levels of Aβ. As a consequence, they develop amyloid pathology that is similar to that found in brains from patients with Alzheimer’s disease^22^. Significant Aβ_42_ levels are detected from 1 month of age in this model, and the first amyloid plaques are detectable in the hippocampus at around 3-4 months of age^22^. This progression can be related to early and intermediate stages of human Alzheimer’s disease. Spontaneous excitatory post-synaptic currents (sEPSCs) were recorded for CA1 pyramidal neurons by whole-cell patch-clamp, using samples from 1-, 2-, and 3-month-old mice. An increase in sEPSC frequency for CA1 neurons was observed in 1-month-old APP/PS1 mice (0.20 ± 0.06 Hz in APP/PS1 mice *versus* 0.08 ± 0.03 Hz in WT; *p* = 0.0266; Fig. 1A, B). This hyperactivity gradually shifted to hypoactivity in 3-month-old APP/PS1-21 mice (0.03 ± 0.01 Hz in APP/PS1 mice *vs* 0.08 ± 0.02 Hz in WT; *p* = 0.0029; Fig. 1A, B). At 2 months old, we observed a transitional situation, with both hyperactive and hypoactive neurons (Fig. 1B). The sEPSC amplitude was unaffected in 1-month-old animals (33.7 ± 1.9 pA in APP/PS1 mice *vs* 33.0 ± 2.3 pA in WT; *p* = 0.9551; Fig. 1A, B) and was reduced at 3 months of age (28.9 ± 1.0 pA in APP/PS1 mice *vs* 35.2 ± 1.9 pA in WT; *p* = 0.0038; Fig. 1A, B). This decrease in amplitude exacerbated the CA1 neuronal hypoactivity.

**Figure 1.**
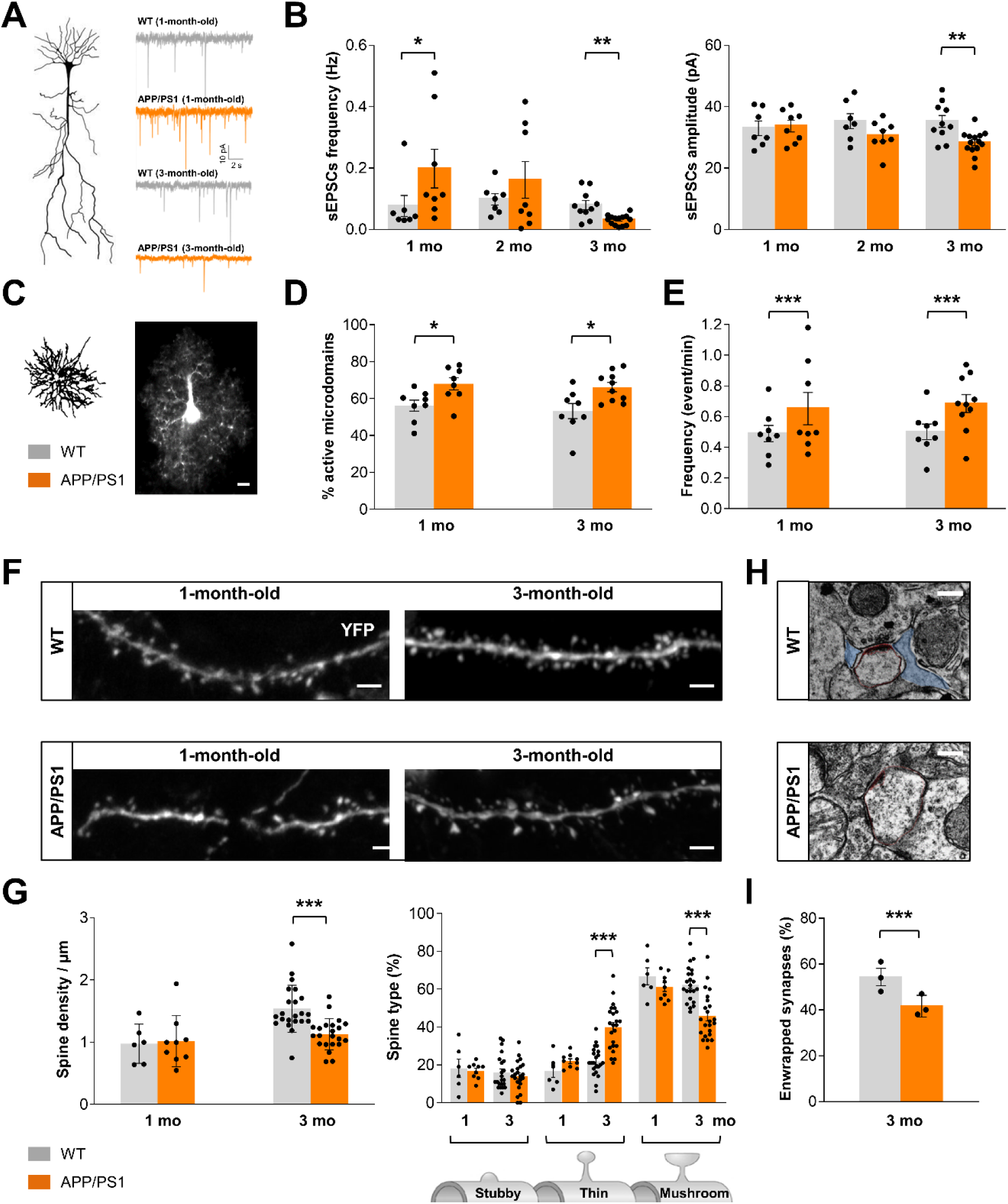
Time course of early functional and structural effects on neurons and astrocytes in APP/PS1-21 mice. **(A)** Representative traces from voltage-clamp recordings for CA1 pyramidal cells from 1- and 3-month-old APP/PS1-21 mice (orange) and their age-matched WT littermates (grey). Cells were held at -60 mV. **(B)** Histograms showing spontaneous EPSC frequency and amplitude measured for CA1 pyramidal neurons from APP/PS1-21 mice (orange) and WT littermates (grey) at 1 month old (*n* = 8 neurons from 6 animals in APP/PS1 and 7 neurons from 5 animals in WT), 2 months old (*n* = 8 neurons form 5 animals in APP/PS1 and 7 neurons from 4 animals in WT), and 3 months old (*n* = 14 neurons from 7 animals in APP/PS1 and 10 neurons from 5 animals in WT). **(C)** A single patch-clamp Fluo-4-loaded astrocyte in the *stratum radiatum* from a WT mouse. Fluorescence variations within the astrocytic arbour were analysed in 1-µm^2^ subregions. Scale bar: 5 µm. **(D)** Proportion of astrocyte subregions displaying calcium activity in APP/PS1-21 (orange) and WT littermates (grey) at 1 month old (*n* = 8 astrocytes from 7 animals in APP/PS1 and 8 astrocytes from 6 animals in WT) and 3 months old (*n* = 10 astrocytes from 4 animals in APP/PS1 and 8 astrocytes from 3 animals in WT). **(E)** Frequency of calcium events in active microdomains in APP/PS1-21 (orange) and WT littermates (grey) at 1 month old (*n* = 650 microdomains from 8 astrocytes in APP/PS1 and 454 microdomains from 8 astrocytes in WT) and 3 months old (*n* = 1504 microdomains from 10 astrocytes in APP/PS1 and 840 microdomains from 8 astrocytes in WT). Each dot corresponds to the mean frequency recorded in individual astrocytes. **(F)** Representative images of regions of CA1 pyramidal neuron dendrites expressing eYFP in Thy1-eYFP-H-APP/PS1-21 mice and their WT littermates at 1 month old (left) and 3 months old (right). **(G)** Quantification of spine density and morphology in APP/PS1-21 (orange) and their WT littermates (grey) at 1 month old (*n* = 9 dendrites from 2 animals in APP/PS1 and 6 dendrites from 2 animals in WT) and 3 months old (*n* = 23 dendrites from 3 animals in APP/PS1 and 22 dendrites from 3 animals in WT). Scale bar = 2 µm. **(H)** Electron micrographs of *stratum radiatum* from 3-month-old WT and APP/PS1-21 mice showing synapse enwrapping by astrocytic perisynaptic processes in WT (blue) and the lack thereof in APP/PS1-21. Approximate scale bar: 0.5 μm. **(I)** Quantification of the proportion of enwrapped synapses in 3-month-old WT (n = 441 synapses from 3 mice) and APP/PS1-21 mice (n = 480 synapses from 3 mice). Each dot corresponds to the mean proportion measured in each individual mouse. *, *p* < 0.05; **, *p* < 0.01; ***, *p* < 0.001 (Mann-Whitney or Fischer’s exact test).

To investigate astrocytic calcium activity, individual astrocytes were loaded with Fluo-4 dye to detect the area covered by single-cell processes^12, 24^ (Fig. 1C). Within these processes, the proportion of active calcium microdomains (≍ 1 µm^2^) was observed to be increased in 1-month-old APP/PS1 mice relative to their WT littermates (67.9 ± 3.3% in APP/PS1 mice *vs* 56.2 ± 2.9% in WT; *p* = 0.0145; Fig. 1D) and remained high in 3-month-old mice (66.2 ± 2.5% in APP/PS1-21 mice *vs* 53.3 ± 4.1% in WT; *p* = 0.0266; Fig. 1D). The frequency of these calcium events within each active microdomain was increased at 1 month of age in APP/PS1-21 mice (0.65 ± 0.10 events/min in APP/PS1-21 mice *vs* 0.49 ± 0.05 events/min in WT; *p* < 0.001; Fig. 1E) and remained high at 3 months old (0.68 ± 0.06 events/min in APP/PS1-21 mice *vs* 0.50 ± 0.05 events/min in WT; *p* < 0.001; Fig. 1E). Thus, astrocyte calcium hyperactivity was stable over time. Based on this result, it appears that Aβ overproduction has an early impact on both astrocytic and neuronal activity, triggering sustained astrocyte hyperactivity in 1- and 3-month-old mice, whereas the temporary neuronal hyperactivity recorded at 1 month shifted to hypoactivity at 3 months old.

We next examined whether neuronal activity disruption was associated to structural modifications. We analysed the spine density and morphology of pyramidal CA1 neurons in transgenic Thy1-eYFP-H mice^23^ cross-bred with APP/PS1-21 mice (Fig. 1F). At 1 month, there was no difference in either spine density or morphology in APP/PS1-21 *versus* WT mice (Fig. 1G), whereas at 3 months old, spine density was reduced in APP/PS1-21 animals (1.13 ± 0.05 spines/µm) relative to WT controls (1.54 ± 0.08 spines/µm; *p* < 0.001; Fig. 1F, G). Concomitantly, the proportion of immature thin spines increased (40 ± 2.5%) in APP/PS1-21 compared to WT (21.9 ± 1.6%; *p* < 0.001; Fig. 1G) and the proportion of mature mushroom spines was reduced at 3 months old (45.7 ± 2.5% in APP/PS1-21 *vs* 62.4 ± 2.0% in WT; *p* < 0.001; Fig. 1G). The astrocytic coverage of *stratum radiatum* synapses was then analysed by electron microscopy (Fig. 1H) and a reduction in the proportion of tripartite synapses was clearly observed in 3-month-old transgenic mice (41.7 ± 2.8% enwrapped synapses in APP/PS1-21 *vs* 54.3 ± 3.7% in WT; *p* < 0.0001; Fig. 1I). Taken together, these data indicate that the decrease in CA1 neuronal sEPSC frequency and amplitude was associated with morphological remodelling of the cells, including a significant reduction in spine density and maturity at 3 months old in APP/PS1-21 mice, together with a reduction in the astrocytic coverage of synapses.

### Early hippocampal neuronal hyperactivity is due to Aβ-induced Ca^2+^-dependent gliotransmitter release

We previously showed that acute application of Aβ oligomers (Aβo) to brain slices induced both astrocytic and neuronal hyperactivity within mouse hippocampus, similar to what was observed in 1-month-old APP/PS1-21 mice^12^. As this early Aβ-induced astrocytic calcium hyperactivity was not due to the increased neuronal activity^12^, we wondered whether, conversely, astrocyte calcium hyperactivity could be involved in establishing Aβo-induced neuronal hyperactivation. To elucidate this question, we selectively blocked calcium dynamics in a single astrocyte in the *stratum radiatum* by intracellular dialysis of the Ca^2+^ chelator BAPTA using a patch pipette. Biocytin was also included in the dialysis solution (Fig. 2A)^30^. sEPSCs were then successively recorded in a neighbouring CA1 pyramidal neuron exposed to 100 nM Aβo or not. As previously described^12^, in basal conditions, a 5-min application of 100 nM Aβo induced a massive increase in sEPSC frequency (244.5 ± 37.8% of baseline in Aβo condition *vs* 107.5 ± 8.7% of baseline in physiological condition; *p* = 0.0041; Fig. 2B). When calcium activity was blocked in the astrocytic syncytium, Aβo application no longer induced an increase in sEPSC frequency (78.0 ± 9.2% of baseline in BAPTA + Aβo condition; *p* = 0.0006 relative to Aβo condition; Fig. 2B). These observations indicated that the early neuronal hyperactivity in the hippocampus is mediated by an Aβo-induced astrocyte calcium signal. Thus, astrocytes play an initial role in sensing Aβ toxicity and generating outputs via Ca^2+^ hyperactivity.

**Figure 2.**
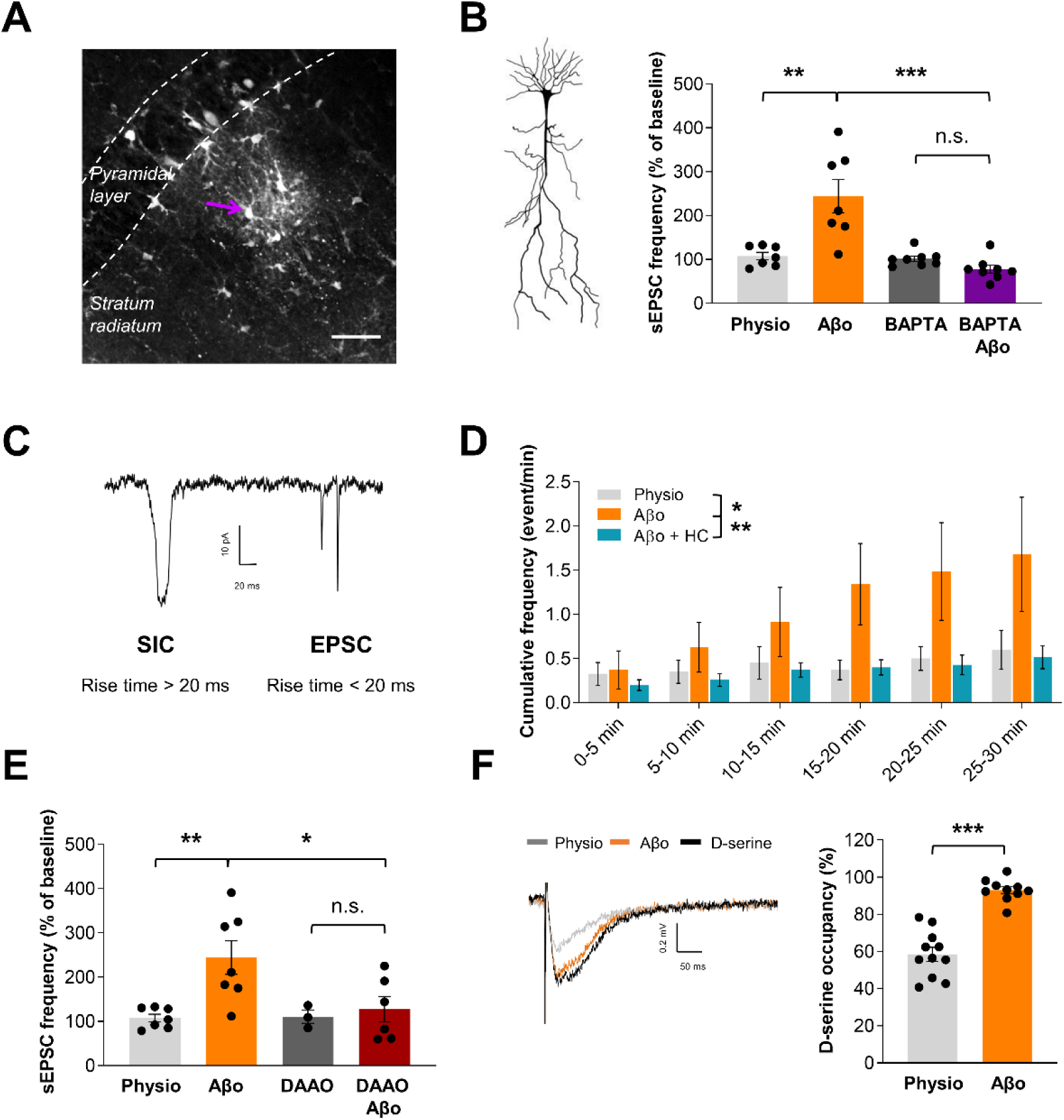
Early Aβo-induced astrocytic calcium hyperactivity triggers glutamate and D-serine release which, in turn, activates nearby neurons. **(A)** Syncytium of astrocytes in CA1 *stratum radiatum* labelled with biocytin after whole-cell diffusion of 10 mM BAPTA in a single astrocyte (purple arrow), delivered through a patch pipette. Scale bar: 50 µm. **(B)** Histogram of CA1 pyramidal neuron sEPSC frequency in physiological conditions (grey; *n* = 7 neurons from 4 mice), 5 min after applying 100 nM Aβo (orange; *n* = 7 neurons from 5 mice), following BAPTA dialysis in the astrocytic syncytium (dark grey; *n* = 8 neurons from 5 mice), or following BAPTA dialysis and application of 100 nM Aβo (purple; *n* = 8 neurons from 5 mice). sEPSC frequency was normalized relative to an initial 5-min recording in physiological conditions. **(C)** Representative electrophysiological trace showing SIC and EPSC recorded for a CA1 pyramidal neuron. **(D)** Time-course of cumulative SIC frequency in physiological conditions (grey; *n* = 8 neurons from 7 mice), following application of 100 nM Aβo to cells (orange; *n* = 7 neurons from 5 mice) or co-application of 100 nM Aβo and 40 µM HC030031 (cyan; *n* = 7 neurons from 5 mice). **(E)** Histogram showing the sEPSC frequency for CA1 pyramidal neurons in physiological conditions (grey; *n* = 7 neurons from 4 mice), 5 min after applying 100 nM Aβo (orange; *n* = 7 neurons from 5 mice), after D-amino acid oxidase (DAAO) treatment in physiological conditions (dark grey; *n* = 3 neurons from 3 mice), or after DAAO treatment followed by application of 100 nM Aβo for 5 min (red; *n* = 6 neurons from 3 mice). **(F)** Representative traces of NMDAR field excitatory post-synaptic potentials (fEPSPs) before (grey), and after (black) application of 10 µM D-serine in control conditions, or 5 min after applying Aβo (orange). **(G)** Histogram showing the NMDAR glycine site occupancy by D-serine (NMDAR fEPSP potentiation) in either physiological conditions (grey; *n* = 11 slices from 4 mice) or 5 min after application of Aβo (orange; *n* = 10 slices from 4 mice). *, *p* < 0.05; **, *p* < 0.01; ***, *p* < 0.001 (Mann-Whitney or Kolmogorov-Smirnov test).

In our previous study, we showed that Aβ-induced astrocytic calcium hyperactivity and subsequent neuronal hyperactivity depend on TRPA1 calcium channel activation^12^. Aβ-induced astrocyte Ca^2+^ hyperactivity is known to trigger glutamate release from astrocytes, this glutamate then activates extrasynaptic NMDAR on nearby neurons^41^. Therefore, to determine whether TRPA1 is involved in this Aβ-dependent astrocytic glutamate release, NMDAR-mediated SICs^28^ were recorded in brain slices (Fig. 2C). In physiological conditions, a low frequency of spontaneous SICs was recorded (0.12 ± 0.05 SICs/min; *n* = 8 neurons), in line with data from the literature^41, 42^. Application of 100 nM Aβo caused the SIC cumulative frequency to increase over a 30-min period (*p* = 0.0260; Fig. 2D). This increase was absent when slices were co-exposed to a TRPA1-specific inhibitor, HC030031^18^ (40 µM), and 100 nM Aβo (*p* = 0.026; Fig. 2D), suggesting that Aβ-induced TRPA1 activation triggers astrocytic glutamate release which then stimulates NMDA receptors on CA1 pyramidal neurons.

The astrocytic TRPA1 calcium channel was shown to be involved in constitutive astrocytic D-serine release within the murine hippocampus, modulating nearby CA1 neuronal function^15^. We therefore investigated whether D-serine was involved in the Aβ-induced neuronal hyperactivity. We first monitored how specific enzymatic D-serine degradation by recombinant DAAO^29^ affected sEPSC frequency. Preincubation with DAAO (0.2 U/ml) consistently resulted in a reduced sEPSC frequency (244.5 ± 37.8% of baseline in Aβo condition *vs* 127.5 ± 28.7% of baseline in DAAO + Aβo; *p* = 0.0411; Fig. 2E). This result suggests that D-serine is involved in Aβ-induced neuronal hyperactivity. We next focused on the occupancy of the NMDAR co-agonist sites controlled by D-serine release^33^. Compared to the saturated condition following application of 10 µM D-serine (*i.e.,* 100% occupancy), in physiological conditions local NMDAR fEPSP amplitude represented 58.5 ± 3.8%. This result indicates that the NMDAR co-agonist sites were not fully occupied at baseline (Fig. 2F). Interestingly, when 100 nM Aβo was applied for 5 min, the local NMDAR fEPSP amplitude reached 93.07 ± 1.8% of the level measured in the saturated condition, indicating that NMDAR co-agonist sites were more saturated in these conditions than at baseline (*p* < 0.0001; Fig. 2F).

Altogether, these data indicated that Aβo has a direct impact on hippocampal astrocytes, triggering compartmentalized calcium hyperactivity^12^ that leads to at least, glutamate and D-serine release. Release of these gliotransmitters elicits subsequent hippocampal neuronal hyperactivity, which is one of the earliest known key events in Alzheimer’s disease pathogenesis^3, 4^. We next examine whether blocking early Aβ-dependent astrocytic calcium hyperactivity could prevent neuronal hyperactivity becoming established. To do so, we performed a longitudinal study in the APP/PS1-21 mouse model to identify the critical juncture at which irreversible progressive neurodegeneration starts in Alzheimer’s disease.

### Chronic inhibition of TRPA1 fully reverses CA1 astrocytic hyperactivity and neuronal hyperactivity in 1-month-old APP/PS1-21 mice

Acute blockade of the TRPA1 channel with HC030031 has been shown to fully restore astrocyte activity to physiological levels in 1-month-old APP/PS1-21 mice, and was also found to totally reverse the early neuronal hyperactivity in this transgenic Alzheimer’s disease mouse model^12^. We therefore investigated whether blocking Aβ-dependent TRPA1 activation as soon as Aβ is overproduced in the brain could protect neurons and maintain neuronal activity. We administered the specific inhibitor of TRPA1 channel (HC030031) intraperitoneally (5 mg/kg body weight) every day from the 14th postnatal day until 1 month of age in APP/PS1-21 mice. As a control, the same volume of vehicle was injected into age-matched and sex-matched control groups, and any effect on the parameters studied in either untreated transgenic or WT littermates was monitored (summarized in Supplementary Table 3).

Results showed that the HC030031 pharmacological treatment prevented the occurrence of hippocampal astrocytic calcium hyperactivity in APP/PS1-21 mice, affecting both the proportion of active microdomains (56.1 ± 3.9% in HC030031-treated mice *vs* 71.8 ± 2.5% in vehicle-treated mice; *p* = 0.0439; Fig. 3A) and the calcium event frequency (0.48 ± 0.02 calcium events/min in HC030031-treated mice *vs* 0.79 ± 0.04 calcium events/min in vehicle-treated mice; *p* < 0.001; Fig. 3B). Thus, these parameters were both restored to physiological values (56.9 ± 3.6% active microdomains and 0.56 ± 0.03 calcium events/min in vehicle-treated WT mice; *p* = 0.7211 and 0.1456, respectively) by HC030031 treatment. Conversely, this compound had no effect on the proportion of active microdomains in WT littermates (53.0 ± 2.7% in HC030031-treated *vs* 56.9 ± 3.6% in vehicle-treated mice; *p* = 0.8627; Fig. 3A) and only slightly decreased the frequency of calcium events within microdomains (0.51 ± 0.02 calcium events/min in HC030031-treated *vs* 0.56 ± 0.03 calcium events/min in vehicle-treated mice; *p* = 0.031; Fig. 3B). Thus, TRPA1 appears to play only a modest role in physiological conditions^12, 15^.

**Figure 3.**
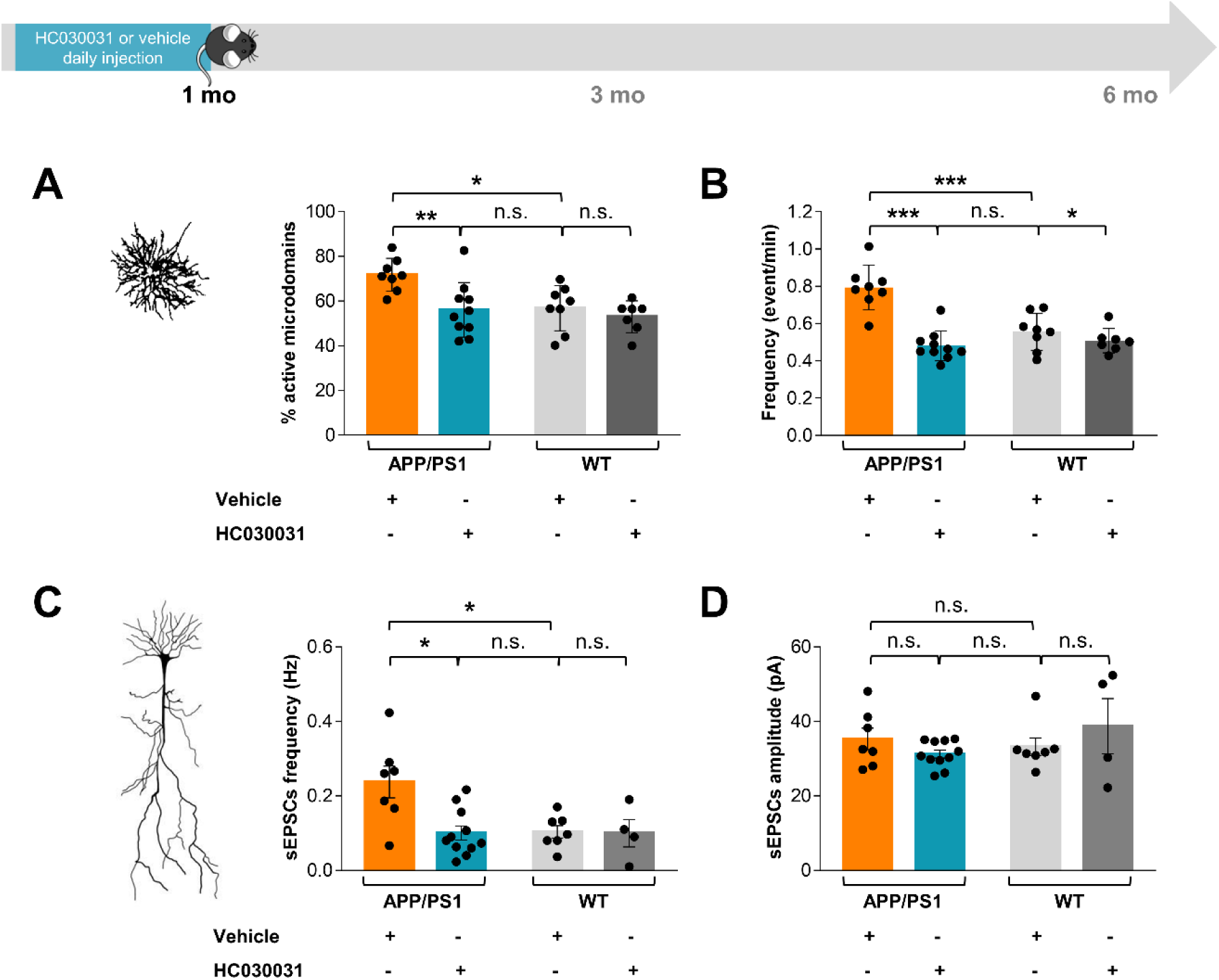
Chronic HC030031 treatment restores neuron and astrocyte activity levels in 1-month-old APP/PS1-21 mice. **(A)** Effect of intra-peritoneal injection of HC030031 or vehicle on the proportion of astrocytic subregions displaying calcium activity in APP/PS1-21 (*n* = 10 astrocytes from 3 HC030031-treated mice and 8 astrocytes from 3 vehicle-treated mice) and WT littermates (*n* = 7 astrocytes from 2 HC030031-treated mice and 8 astrocytes from 3 vehicle-treated mice). **(B)** Effect of intra-peritoneal injection of HC030031 or vehicle on the frequency of calcium activity within astrocytic subregions in APP/PS1-21 (*n* = 771 microdomains from 10 astrocytes in HC030031 condition and 1303 microdomains from 8 astrocytes in vehicle condition) and WT littermates (*n* = 816 microdomains from 7 astrocytes in HC030031 and 995 microdomains from 8 astrocytes in vehicle). **(C, D)** Effect of intra-peritoneal injection of HC030031 or vehicle on sEPSC frequency and amplitude in APP/PS1-21 mice (*n* = 11 neurons from 5 HC030031-treated mice and 7 neurons from 5 vehicle-treated mice) and WT littermates (*n* = 4 neurons from 2 HC030031-treated mice and 7 neurons from 4 vehicle-treated mice). *, *p* < 0.05; **, *p* < 0.01; ***, *p* < 0.001; n.s., not significant (Kruskal-Wallis test).

In addition, HC030031 treatment prevented the occurrence of hippocampal neuronal hyperactivity in transgenic mice (0.10 ± 0.02 Hz following HC030031 treatment *vs* 0.24 ± 0.04 Hz following vehicle treatment; *p* = 0.0311; Fig. 3C), restoring the sEPSC frequency to physiological levels (0.10 ± 0.02 Hz in vehicle-treated WT mice; *p* = 0.7213) at 1 month old. Once again, HC030031 treatment had no effect on WT littermates (0.10 ± 0.04 Hz following HC030031 treatment *vs* 0.10 ± 0.02 Hz following vehicle treatment; *p* > 0.999; Fig. 3C). The sEPSC amplitude was not affected in 1-month-old transgenic mice (Fig. 1B) and HC030031 treatment had no effect on this parameter (*p* > 0.999; Fig. 3D). As described following external Aβo application (Fig. 2D), the overproduction of Aβ in mouse brain at this early stage was associated with defects in astrocytic glutamate release and activation of extra-synaptic NMDA receptors. This activation can be monitored based on the increase in cumulative SIC frequency recorded for CA1 pyramidal neurons in vehicle-treated transgenic mice (*p* = 0.0022; Supplementary Fig. 1A). Interestingly, chronic HC030031 treatment prevented this extra-astrocytic glutamate release in 1-month-old transgenic mice (*p* = 0.0022; Supplementary Fig. 1A), confirming that TRPA1 channel activation plays a pivotal role in this Aβ-induced astrocytic glutamate release.

These data demonstrate that the early inhibition of the TRPA1 channel in this transgenic Alzheimer’s disease mice model is sufficient to prevent hippocampal neuronal hyperactivity at 1 month old, confirming the major role played by astrocytes in the early stages of Aβ-dependent toxicity.

### Chronic inhibition of TRPA1 fully reverses CA1 astrocyte hyperactivity, neuronal hypoactivity, dendritic spine impairment and defective synapse enwrapping in 3-month-old APP/PS1-21 mice

Neuronal hyperactivity is predicted to be a key feature of early-stage Alzheimer’s disease^4–6^, during which it is expected to trigger a vicious cycle culminating in dysregulation of the whole neuronal network. In our model, this further dysregulation manifested as both neuronal hypoactivity and morphological remodelling of dendritic spines from 3 months old (Fig. 1). We thus wondered whether impeding Aβ-induced TRPA1 activation could protect neurons from entering this vicious cycle. Therefore, the specific inhibitor of TRPA1 channel (HC030031) or the vehicle was administered intraperitoneally (5 mg/kg body weight) every day from the 14th postnatal day until 3 months of age in APP/PS1-21 mice. These experiments revealed that chronic TRPA1 inhibition in APP/PS1-21 mice prevented neuronal hypoactivity, as reflected by the sEPSC frequency (0.10 ± 0.01 Hz in HC030031 *vs* 0.04 ± 0.01 Hz in vehicle-treated mice; *p* = 0.01; Fig. 4A) and its amplitude (33.4 ± 1.4 pA in HC030031 *vs* 28.5 ± 1.2 pA in vehicle-treated mice; *p* = 0.0358; Fig. 4B). Levels of neuronal activity in HC030031-treated APP/PS1-21 mice were thus similar to those recorded for vehicle-treated WT littermates (frequency: 0.11 ± 0.01 Hz and amplitude: 32.2 ± 1.1 pA; *p* = 0.386 and 0.6362, respectively). Chronic HC030031 treatment had no effect on sEPSC frequency or amplitude in WT mice (*p* > 0.999 for both parameters; Fig. 4A, B).

**Figure 4.**
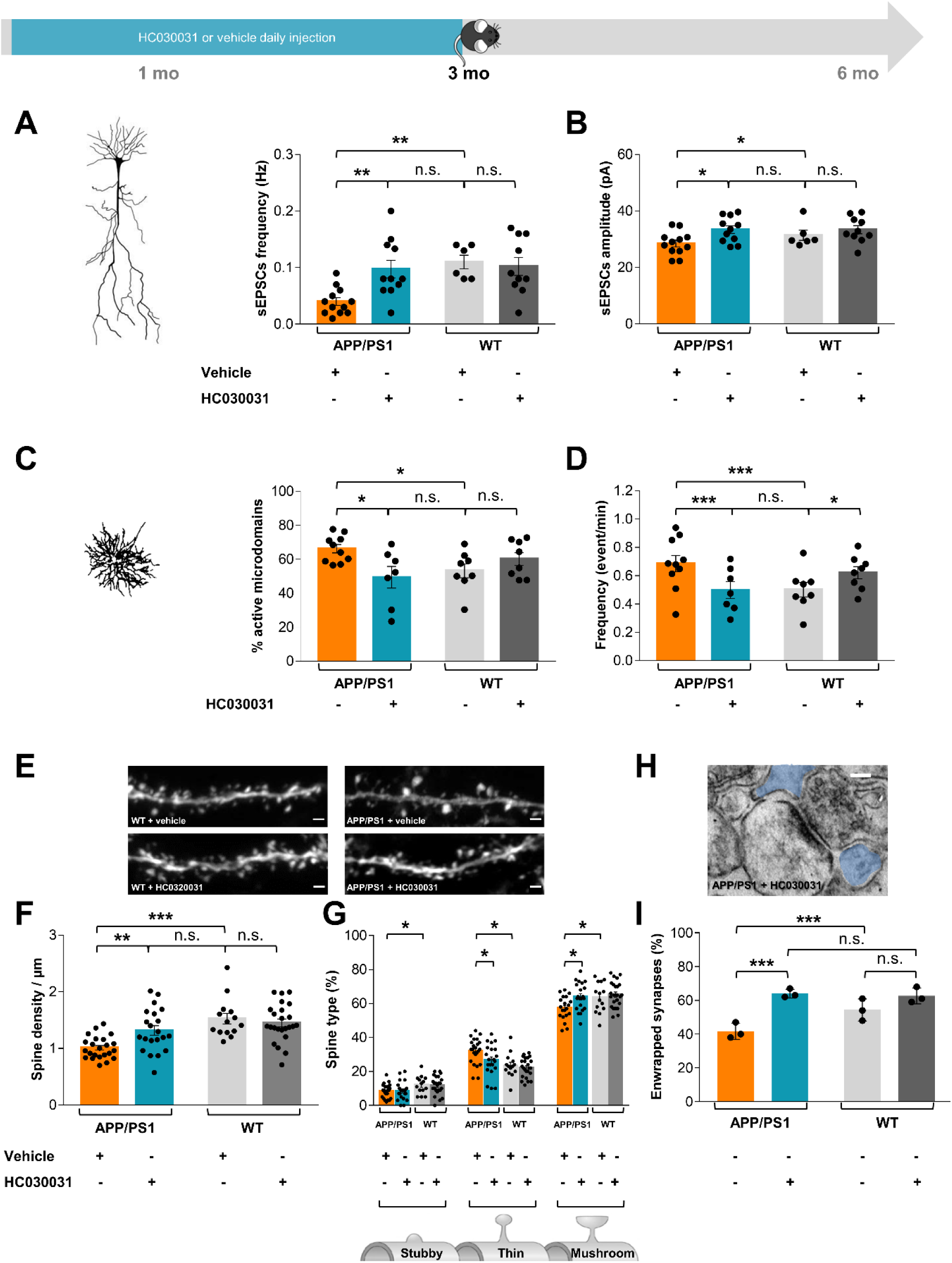
Chronic HC030031 treatment prevents functional and morphological effects on CA1 neurons in 3-month-old APP/PS1-21 mice. **(A, B)** Effect of intra-peritoneal injection of HC030031 or vehicle on sEPSC frequency and amplitude in APP/PS1-21 mice (*n* = 11 neurons from 7 HC030031-treated mice and 12 neurons from 8 vehicle-treated mice) and WT littermates (n = 11 neurons from 6 mice in HC030031 and 10 neurons from 9 mice in vehicle). **(C)** Effect of intra-peritoneal injection of HC030031 on the proportion of astrocytic subregions displaying calcium activity in APP/PS1-21 (*n* = 7 astrocytes from 3 HC030031-treated mice and 10 astrocytes from 3 untreated mice) and WT littermates (*n* = 8 astrocytes from 4 HC030031-treated mice and 8 astrocytes from 3 untreated mice). **(D)** Effect of intra-peritoneal injection of HC030031 on the frequency of calcium activity within astrocytic subregions in APP/PS1-21 (*n* = 1052 microdomains from 7 astrocytes in HC030031-treated condition and 1504 microdomains from 10 astrocytes in untreated condition) and WT littermates (*n* = 1267 microdomains from 8 astrocytes in HC030031-treated condition and 840 microdomains from 8 astrocytes in untreated condition). **(E)** Representative images of areas of CA1 pyramidal neuron dendrites expressing eYFP in Thy1-eYFP-H-APP/PS1-21 mice and their WT littermates following treatment with vehicle or HC030031. **(F)** Effect of intra-peritoneal injection of HC030031 or vehicle on spine density in CA1 neurons in Thy1-eYFP-H-APP/PS1-21 mice (*n* = 21 dendrites from 3 HC030031-treated mice and 14 dendrites from 2 vehicle-treated mice in vehicle) and WT littermates (*n* = 24 dendrites from 3 HC030031-treated mice and 14 dendrites from 2 vehicle-treated mice). **(G)** Effect of intra-peritoneal injection of HC030031 or vehicle on spine typology in CA1 neurons from Thy1-eYFP-H-APP/PS1-21 mice. Spines were classed as thin, stubby, or mushroom. **(H)** Electron micrograph of *stratum radiatum* from a 3-month-old APP/PS1-21 mouse treated with HC030031, showing an example of a synapse enwrapped by astrocytic perisynaptic processes (blue). Approximate scale bar: 0.5 μm. **(I)** Effect of intra-peritoneal injection of HC030031 on the proportion of enwrapped synapses in 3-month-old WT (*n* = 441 synapses from 3 mice in untreated condition and 124 synapses from 3 mice in HC030031-treated condition) and APP/PS1-21 mice (*n* = 480 synapses from 3 mice in untreated condition and 122 synapses from 3 mice in HC030031-treated condition). *, *p* < 0.05; **, *p* < 0.01; ***, *p* < 0.001; n.s., not significant (Kruskal-Wallis or Fischer’s exact test).

Inhibition of the astrocyte calcium hyperactivity observed in 1-month-old mice was also observed at 3 months old after chronic HC030031 treatment of APP/PS1-21 mice. Indeed, both the proportion of active microdomains (49.4 ± 6.4% in HC030031-treated mice vs 66.2 ± 2.5% in untreated mice; *p* = 0.033; Fig. 4C) and the frequency of calcium events within these microdomains (0.50 ± 0.06 events/min in HC030031-treated vs 0.68 ± 0.06 events/min in untreated mice; *p* < 0.0001; Fig. 4D) were reduced, almost reaching physiological levels. Once again, chronic HC030031 treatment had no significant effect on the proportion of active microdomains in WT littermates (60.2 ± 3.8% in HC030031-treated *vs* 53.2 ± 4.1% in untreated mice; *p* = 0.2786; Fig. 4C) although a slight increase in the frequency of calcium events within these microdomains was observed (0.62 ± 0.04 calcium events/min in HC030031-treated *vs* 0.50 ± 0.05 calcium events/min in untreated mice; *p* = 0.001; Fig. 4D).

To determine how this chronic treatment affected spine density and morphology, we administered HC030031 or vehicle from the 14th postnatal day until 3 months of age in Thy1-eYFP-H-APP/PS1-21 transgenic mice. Chronic TRPA1 inhibition led to normalization of spine density (1.32 ± 0.08 spines/µm in HC030031-treated mice *vs* 1.01 ± 0.04 spines/µm in vehicle-treated mice; *p* = 0.0019; Fig. 4E, F) and spine maturation (26.7 ± 2.0% thin spines in HC030031-treated mice *vs* 32.0 ± 1.6% thin spines in vehicle-treated mice and 64.0 ± 1.7% mushroom spines in HC030031-treated mice *vs* 57.3 ± 1.5% mushroom spines in vehicle-treated mice; *p* = 0.05 and 0.0117, respectively; Fig. 4G).

In addition, electron microscopy analysis revealed that the astrocytic coverage of *stratum radiatum* synapses was fully restored in HC030031-treated transgenic mice (64.0 ± 1.5% enwrapped synapses in HC030031-treated APP/PS1-21 mice *vs* 41.7 ± 2.7% in untreated APP/PS1-21 mice; *p* < 0.0001; Fig. 4H, I). In WT mice, in contrast, HC030031 treatment had no effect on astrocyte coverage (62.7 ± 2.7% enwrapped synapses in HC030031-treated WT mice vs 54.3 ± 3.7% in untreated WT mice; *p* = 0.0813; Fig. 4H, I). These structural alterations were associated with restored neuron-astrocyte interactions, as demonstrated by functional study of how astrocytic glutamate release influenced nearby neurons. In 3-month-old APP/PS1 mice, a strong decrease in SIC cumulative frequency was observed in vehicle-treated transgenic mice compared to WT littermates (*p* = 0.026; Supplementary Fig. 1B), which is consistent with the unwrapping of *stratum radiatum* synapses (Fig. 4I). Chronic HC030031 treatment increased the cumulative frequency of SICs (p = 0.0022; Supplementary Fig. 1B) to a level equivalent to that recorded for WT mice (*p* = 0.474), suggesting functional normalization of the interaction between astrocytes and neighbouring neurons.

Overall, these data showed that early chronic inhibition of TRPA1 channels is sufficient to prevent neuronal hyperactivity becoming established. This hyperactivity appears to be the driving force for early progressive failures, leading to the loss of functional dendritic spines and subsequent neuronal hypoactivity. Simultaneously, this chronic inhibition of TRPA1 channels restored the proportion of tripartite synapses in the *stratum radiatum*, depletion of which is likely to be significantly involved in the complex noxious Aβ-induced process.

### Chronic inhibition of TRPA1 promotes amyloid fibril compaction in plaques and strengthens the astroglial plaque barrier in 6-month-old APP/PS1-21 mice

Accumulation of amyloid-β deposits is a characteristic feature of Alzheimer’s disease that develops late (*i.e.* from 4-month-old) in the hippocampus of APP/PS1-21 mice^22^. We thus determined how daily HC030031 treatment from the beginning of Aβ overproduction affected Aβ plaque formation in 6-month-old APP/PS1-21 mice. Chronic treatment with HC030031 had no effect on hippocampal Aβ plaque numbers - as measured by thioflavin S staining (2.1 ± 0.3 plaques/mm^2^ in vehicle-treated mice *vs* 1.8 ± 0.4 plaques/mm^2^ in HC030031-treated mice; *n* = 3 mice for each condition; *p* = 0.2849; Supplementary Fig. 2A). However, the plaques in treated and untreated animals were clearly visually distinct (Fig. 5A). We therefore analysed both surface area and mean plaque intensity as indicators of plaque compaction^38^, by co-labelling thioflavin S and amyloid fibrils (using OC antibodies)^36^. Diffuse amyloid fibril-staining was observed around thioflavin S-labelled plaques in vehicle-treated transgenic mice, whereas staining was more compact in HC030031-treated transgenic mice (Fig. 5A). The area of Aβ plaques was reduced both in terms of thioflavin S labelling (281.5 ± 39.9 µm^2^ in vehicle-treated mice *vs* 152.6 ± 16.2 µm^2^ in HC030031-treated mice; *p* = 0.0024; Fig. 5A, B) and OC staining (179.1 ± 93.6 µm^2^ in vehicle-treated mice *vs* 100.2 ± 64.0 µm^2^ in HC030031-treated mice; *p* = 0.003; Fig. 5A, B). In HC030031-treated transgenic mice the mean intensity for OC staining was increased (24.1 ± 2.1 in vehicle-treated mice vs 46.2 ± 8.0 in HC030031-treated mice; *p* = 0.0311; Fig. 5A, B). This increase specifically reflects amyloid fibril compaction in amyloid plaques^38^. This result was confirmed by dot-blot measurements, where the relative intensity of OC staining increased for samples from HC030031-treated transgenic mice (48.4 ± 2.2 in HC030031-treated mice *vs* 33.3 ± 2.6 in vehicle-treated transgenic mice; *p* = 0.0047; Supplementary Fig. 2A, B). Taken together, these data reflect a reorganization of Aβ deposits, with increased Aβ fibril assembly, and greater compaction of these Aβ fibrils^38, 43^ in plaques in HC030031-treated mice. Sequestration of Aβ in plaques is known to contain their toxic effects, rendering them functionally innocuous^44^.

**Figure 5.**
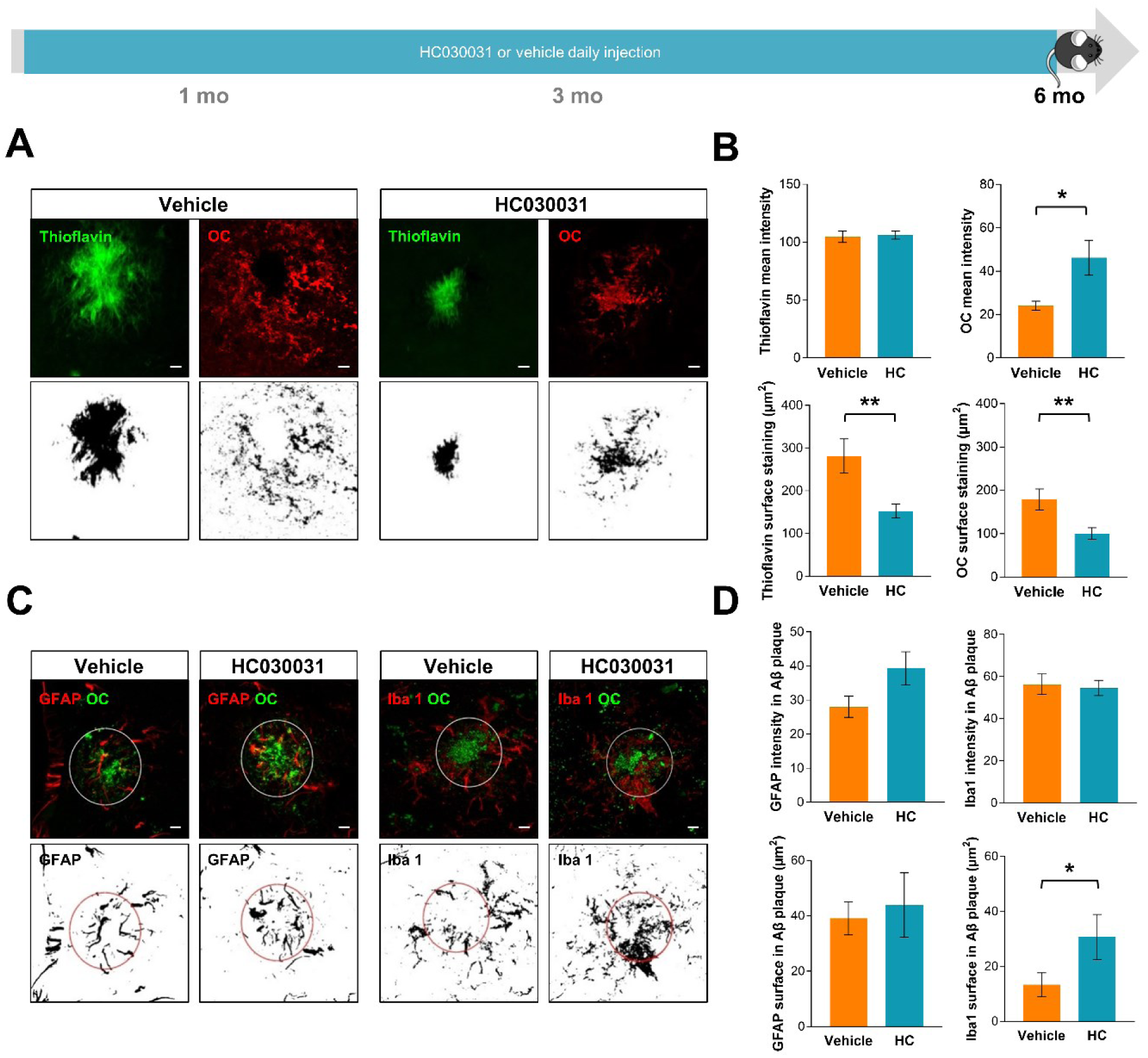
Chronic HC030031 treatment increases plaque compaction and astroglial plaque barrier development in 6-month-old APP/PS1-21 mice. **(A)** Representative examples of high-resolution confocal images of hippocampal plaques co-stained for Thioflavin S (green) and amyloid fibrils (OC antibodies; red). Hippocampal sections from 6-month-old APP/PS1-21 mice treated daily with vehicle or HC030031. Corresponding threshold images used to quantify stained area and mean fluorescence intensity are also shown. Scale bar = 5 µm. **(B)** Quantification of Thioflavin S and amyloid fibril staining area and mean fluorescence intensity in the hippocampus of vehicle-treated (orange; *n* = 15 plaques from 3 mice) and HC030031-treated (cyan; *n* = 22 plaques from 3 mice) APP/PS1-21 mice. **(C)** Representative examples of high-resolution confocal images of hippocampus plaque-associated astrocytes (stained with GFAP, red; and OC, green) and microglia (stained with Iba1, red; and OC, green) in vehicle-treated or HC030031-treated APP/PS1-21 mice. The white ring delimits the amyloid plaque area. Bottom panels show corresponding threshold images used to quantify the stained area and determine the mean fluorescence intensity within the plaque (red ring). Scale bar = 5 µm. **(D)** Quantification of GFAP and Iba 1 staining intensity and area in hippocampal amyloid plaques from vehicle-treated (orange; *n* = 15 plaques from 3 hippocampi) and HC030031-treated (cyan; *n* = 16 plaques from 3 hippocampi) APP/PS1-21 mice. *, *p* < 0.05; **, *p* < 0.01; ***, *p* < 0.001; n.s., not significant (Mann-Whitney test).

Astrocytes and microglia are frequently associated with plaques in Alzheimer’s disease and form a dense anatomical barrier around Aβ plaques which limits the diffusion of neurotoxic Aβ^38, 45^. We therefore studied the distribution of and morphological changes to these cells in the periplaque zone in the hippocampus by immunostaining for GFAP and Iba-1. In vehicle-treated APP/PS1-21 mice, microglia and astrocytes with enlarged cell bodies and processes were observed. These morphologies are consistent with changes associated with reactive microgliosis and astrogliosis reported for this mouse model^22^. GFAP-positive astrocytes and Iba-1-positive microglia were clustered around plaques in the hippocampus from APP/PS1-21 mice, but appeared to invade the plaque core in HC030031-treated mice (Fig. 5C). We quantified the stained area and mean intensity of GFAP-positive and Iba-1-positive processes surrounding OC-positive Aβ plaques using high-resolution Z-stack confocal microscopy imaging^39^. No difference in GFAP staining area was found within Aβ plaques in HC030031-treated transgenic mice (43.9 ± 11.6 µm^2^ in HC-treated mice *vs* 39.1 ± 6.0 µm^2^ in vehicle-treated mice; *p* = 0.8518; Fig. 5D), but there was a slight increase in mean GFAP intensity in HC030031-treated transgenic mice (39.4 ± 4.8 in HC-treated mice *vs* 28.0 ± 3.1 in vehicle-treated mice; *p* = 0.0893; Fig. 5D). In contrast, a large increase in the area stained with Iba-1 was detected (30.8 ± 8.2 in HC030031-treated mice *vs* 13.3 ± 4.3 in vehicle-treated mice; *p* = 0.0330; Fig. 5D), once again without difference in mean intensity for this marker (54.6 ± 3.6 in HC-treated mice *vs* 56.4 ± 4.8 in vehicle-treated mice; *p* = 0.3791; Fig. 5D). These results suggest that chronic HC030031 treatment may improve the microglial response to plaques, by forming a barrier that could help to contain the diffusion of neurotoxic Aβ fibrils^38^.

Overall, these data revealed that blocking TRPA1 from the beginning of Aβ overproduction has an effect on the compaction of plaques and the reaction and clustering of glia around plaques. The results highlight the reciprocal synergistic interaction between astrocytes and microglia, and confirm that microglial processes, more than astrocyte processes, are a critical determinant in the degree of amyloid compaction achieved. This compaction prevents plaque expansion and the escape of neurotoxic protofibrillar Aβ into the area surrounding plaques^38^.

### Chronic inhibition of TRPA1 prevents spatial working-memory defects in 6-month-old APP/PS1-21 mice

Having demonstrated extensive cellular and morphological effects of TRPA1-inhibition in our mouse model of Alzheimer’s disease, we wished to investigate its effects on cognitive parameters of the disease. Spatial working-memory-based tasks are extensively used to test murine memory; these tasks closely model aspects of the working-memory deficit observed in Alzheimer’s disease^46^. Here, we wished to determine whether inhibiting Aβ toxicity through TRPA1 blockade could prevent the occurrence of such Alzheimer’s disease-related memory defects. A previous study showed that genetic ablation of TRPA1 in APP/PS1-21 mice restored spatial learning and memory function in these mice^47^, thus flagging the TRPA1 channel as a crucial player in Alzheimer’s disease pathogenesis and consequently a potential therapeutic target. Once again, we used HC030031 (5 mg/kg body weight) to specifically inhibit the TRPA1 channel and compared its effects to those of vehicle. Compounds were administered intraperitoneally to APP/PS1-21 and WT mice every day from the 14th postnatal day until 6 months of age before testing their behaviour. The effect of vehicle in either transgenic mice or their WT littermates are summarized in Supplementary Table 3. The performance of 6-month-old mice was tested for hippocampus-dependent spatial working- and reference-memory tasks based on the Barnes maze test paradigm^43^. Over the 3-day acquisition period (8 trials), the time required to reach the escape box (escape latency) decreased for both transgenic and WT mice as they became familiar with the setup, indicating that both groups of animals learned to use spatial cues to find the escape box (Fig. 6B). However, APP/PS1-21 mice consistently reached the escape box more slowly than controls during this learning phase, suggesting a spatial working-memory defect in these transgenic mice (measured based on the area under the curve for escape latency: 731.8 ± 91.1 in vehicle-treated APP/PS1-21 mice *versus* 554.1 ± 78.9 in vehicle-treated WT mice; *p* = 0.0012; Fig. 6A, C). Motor function, based on the speed at which mice moved, measured across all days, was similar for all groups (*p* = 0.1718; Fig. 6D). To assess reference memories formed as a result of the repeated trials, during which the task parameters remained unchanged, mice were subjected to a probe test in which the escape box was removed. In this test, the number of explorations of the target area was lower in APP/PS1-21 mice (12.8 ± 1.0 in vehicle-treated animals) compared to WT (19.3 ± 1.5 in vehicle-treated individuals; *p* = 0.0374; Fig. 6E), suggesting that spatial information was not fully incorporated into reference memory in the transgenic mice. Based on these results, we concluded that both working- and reference-spatial memories were impaired in 6-month-old APP/PS1-21 mice compared to their WT littermates. Treatment with HC030031 from the beginning of Aβ overproduction partly alleviated these mnesic defects. Indeed, working-memory defects identified during the learning phase in transgenic mice were fully compensated (AUC: 595.9 ± 77.2 in HC030031-treated APP/PS1-21 mice *versus* 731.8 ± 91.1 in vehicle-treated APP/PS1-21 mice; *p* = 0.0089; Fig. 6B and C), with performance becoming similar to WT (554.1 ± 78.9; *p* = 0.8054). Conversely, long-term HC030031 treatment did not improve reference memory, since the number of explorations of the target area during the probe test was lower in HC030031- and vehicle-treated transgenic mice than in WT mice (10.7 ± 1.0 in HC030031-treated APP/PS1-21 mice *versus* 12.8 ± 1.0 in vehicle-treated APP/PS1-21 mice; *p* = 0.5354; Fig. 6E). Reassuringly, chronic HC030031 treatment had no effect on working- or reference-memory performance in WT mice (Fig. 6B-E). Thus, these data revealed that blocking TRPA1 channel in early preclinical stage restored spatial learning and working-memory deficit, classical outcome of Alzheimer’s disease progression.

**Figure 6.**
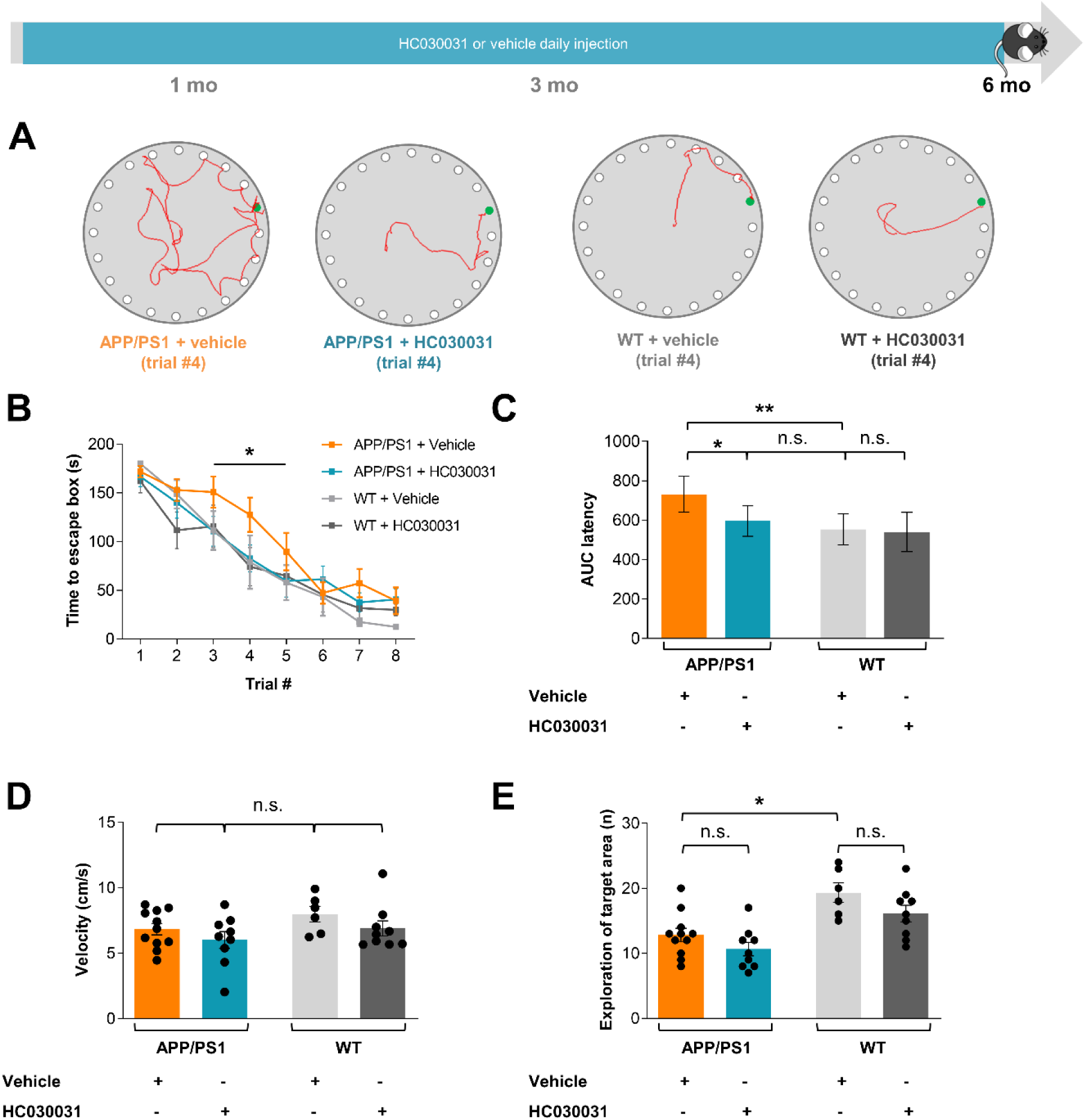
Chronic HC030031 treatment partially prevents spatial memory deficits in 6-month-old APP/PS1-21 mice. **(A)** Representative examples of search strategies developed for each group of mice in the Barnes maze test used as a spatial-learning task. Mice had to locate an escape box (green hole) in a mildly aversive environment. Mice were treated daily for 6 months with vehicle or HC030031. **(B)** Learning curve based on escape latency for vehicle-treated WT (light grey; *n* = 6 mice), vehicle-treated APP/PS1-21 (orange; *n* = 10 mice), HC030031-treated WT (dark grey; *n* = 9 mice), and HC030031-treated APP/PS1-21 (cyan; *n* = 11 mice) mice. The learning curve was similar for all groups except for vehicle-treated APP/PS1-21 mice, for which longer escape latencies were recorded in trials 3, 4, and 5. **(C)** Area under the curve (AUC) for latency to reach the escape box for APP/PS1-21 + vehicle (orange) or HC030031-treated mice (cyan). **(D)** Mouse movement velocity in the Barnes maze across all days. **(E)** Probe test for vehicle-treated APP/PS1 mice (orange) compared to vehicle-treated WT mice (light grey); and following HC030031-treatment in APP/PS1-21 (cyan) or WT mice (dark grey). *, *p* < 0.05; **, *p* < 0.01; ***, *p* < 0.001; n.s., not significant (Kruskal-Wallis or 2-way ANOVA test).

## Discussion

Aberrant hippocampal astrocytic and neuronal hyperactivity appears to be an early causal factor in synaptic remodelling leading to cognitive decline in Alzheimer’s disease. This destabilized hippocampal network is evident in both animal models of Alzheimer’s disease^4^ and in presymptomatic Alzheimer’s disease patients, where it heralds and contributes to the future decline^5, 48, 49^. Our previous results indicated that Aβ-dependent astrocyte activation through the TRPA1 channel accounts for nearby neuronal hyperactivity in the mouse hippocampus during the early stages of Alzheimer’s disease^12^. Here, we showed that astrocytic calcium hyperactivity triggers glutamate and D-serine release, both of which are involved in inducing neuronal hyperactivity. We next investigated the contribution of astrocytes to early neuronal remodelling, and its deleterious consequences, by blocking TRPA1 channel activation from the onset of Aβ overproduction in APP/PS1-21 mice. Early treatment with TRPA1 inhibitor (HC030031) normalized astrocyte activity, avoided loss of synapse enwrapping, and protected structural and functional neuronal integrity. In the longer term, TRPA1 inhibition strengthened the anatomical astroglial barrier around Aβ plaques and promoted Aβ fibrils compaction. This early multi-level neuroprotection appears to prevent subsequent declines in spatial working-memory. Overall, the data presented in this article highlight the major role played by astrocytes in the initiation of a pathological context through their regulation of neuronal structure and function. Blockade of this critical event appears to be sufficient to avoid deleterious collateral damage (Fig. 7).

**Figure 7.**
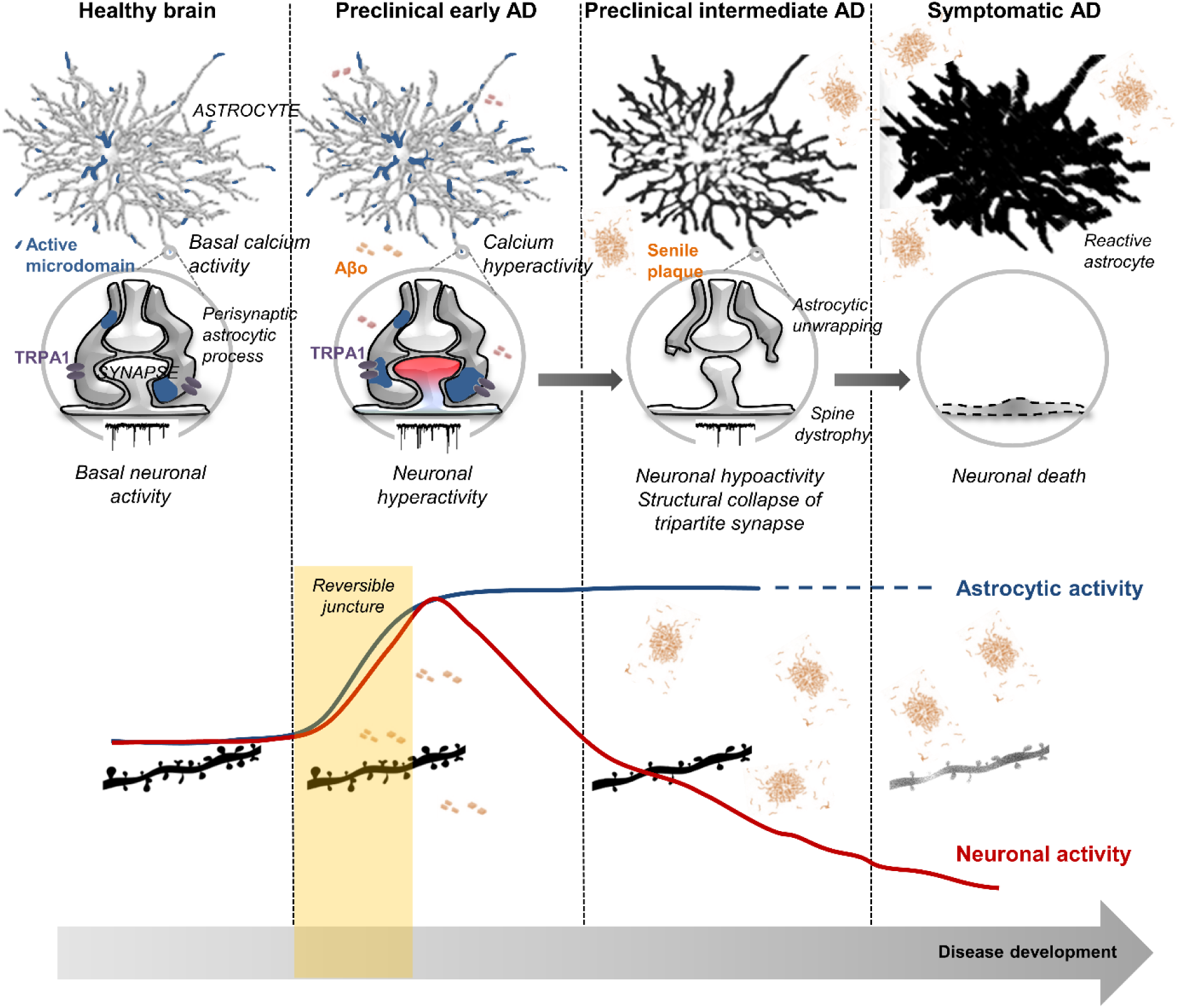
Potential involvement of astrocytes in regulating synaptic activity during the early stages of Alzheimer’s disease and progression towards irreversible critical junctures. In physiological conditions, astrocytes exhibit spontaneous calcium activity that contributes to maintaining the structural and functional integrity of the synapse (active microdomains in blue). In the early stages of Alzheimer’s disease, the presence of oligomeric amyloid-β peptide (Aβo) disrupts the physiological functioning of the astrocyte. Through an activating effect on the calcium channel TRPA1, the astrocyte becomes hyperactive and releases gliotransmitters which induce hyperactivity in neighbouring neurons. These phenomena combine to produce synaptic toxicity (neuronal hypoactivity, morphological collapse of dendritic spines, astrocytic unwrapping, etc.) and neuroinflammation, preludes to memory disorders and neuronal death (symptomatic Alzheimer’s disease). Blockade of the initial events with TRPA1 inhibitor seems to be sufficient to prevent the vicious cycle of Alzheimer’s disease neurodegeneration.

Recently, loss of cellular homeostasis was hypothesized to trigger the start of the clinical Alzheimer’s disease phase, whereas Aβ plaques and phosphorylated Tau accumulations are considered downstream features of the disease^50^. Our data confirmed that a failure in astrocyte-neuron interplay underlies the firing instability in neuronal circuits and constitutes the common drivers of Alzheimer’s disease pathogenesis^51^. In the Alzheimer’s disease transgenic mouse model used here, we discovered a critical time window starting from Aβ overproduction which then causes hyperactivity in hippocampal astrocytes and neurons. These early functional defects trigger temporal sequences of undefined dyshomeostatic events leading to synaptic impairment and neuronal hypoactivity. Many researchers speculate that new therapeutic strategies will need to control this early dysregulation to prevent the collateral damage caused by neuronal hyperactivity, and therefore protect neural networks^4, 5, 51, 52^.

Despite their probable involvement in these complex and critical cellular phases of Alzheimer’s disease^50^, astrocyte behaviour and reactions during the early phases of Alzheimer’s disease remain largely undefined. The results presented here clearly showed, for the first time, that astrocytic Ca^2+^ hyperactivity is directly linked to the subsequent Aβ-dependent neuronal hyperactivity, since blocking calcium dynamics in the astrocyte network by intracellular dialysis of BAPTA fully prevented the neuronal effects of Aβ. We further showed that Aβ affects astrocytes by triggering Ca^2+^-dependent release of neuroactive substances, such as glutamate and D-serine, that will subsequently influence synaptic and extra-synaptic transmission^24, 31^. Augmenting D-serine availability at glutamatergic synapses will increase the amount of activable NMDAR^29^ and, in complement, astrocytic glutamate release will activate extrasynaptic NMDAR^41, 53^. These two neuro-transmitters will thus synergistically increase synaptic activity in pyramidal neurons, probably initiating Aβ-induced synaptotoxicity leading to the disruption of neuronal signalling and loss of synapses^41, 54, 55^.

In our Alzheimer’s disease transgenic mouse model, the compartmentalized astrocyte calcium signalling hyperactivity and resulting hyperactivity of nearby CA1 neurons were both detectable from the onset of Aβ overproduction, *i.e.*, in 1-month-old mice^12^. These early functional alterations evolved progressively and were accompanied by morphological remodelling of astrocytes and neurons. Indeed, the early neuronal hyperactivity observed in 1-month-old transgenic mice gradually led to hypoactivity in 3-month-old APP/PS1-21 mice. High-resolution studies showed that morphological changes, such as a decrease in dendritic spine density and shrinkage of dendritic spines, were present in 3-month-old APP/PS1-21 mice, whereas they were not noticeable during the early hyperactive stage. These alterations occurred alongside an extensive and significant reduction in the enwrapping of synapses by astrocytes in the *stratum radiatum* in 3-month-old transgenic animals, as revealed by ultrastructural electron microscopy studies. These structural changes profoundly altered neuron-astrocyte interactions, and together with the decrease in dendritic spine density, the alterations to synapse enwrapping could emphasize CA1 neuronal dysfunction and glutamate dyshomeostasis^56^. These structural and functional repercussions appeared to progressively transform, becoming irreversible phenomena.

Within Alzheimer’s disease pathogenesis, the initial astrocytic/neuronal hyperactivity could thus be considered a critical juncture that will progressively lead to irreversible cellular effects^3, 4, 50^. Our previous data suggested that the TRPA1 channel is at the forefront of Aβ-induced astrocyte hyperactivity and its neuronal consequences^12^. In the present longitudinal study, we found that prophylactic treatment with a TRPA1 inhibitor normalized initial astrocyte activity, reaching levels equivalent to those detected in healthy WT animals. This normalization prevented the triggering of hippocampal neuronal hyperactivity, confirming a pivotal role for astrocytes in preserving synapse integrity. Interestingly, this result demonstrates that the early neuronal hyperactivity is the starting point for subsequent synaptic dysfunction, structural collapse, and cognitive defects. Blocking it appears to be sufficient to prevent the vicious cycle of Alzheimer’s disease neurodegeneration, and to partly restore spatial memory deficits. In amnestic mild cognitive impairment (aMCI) patients, reducing hippocampal hyperactivity with a low dose of leviracetam - a GABA_A_ receptor agonist - has already proven its effectiveness in improving cognition and reducing progression to the dementia-phase in human Alzheimer’s disease^57^. In rodents, a low dose of diazepam also showed beneficial effects in the early pre-dementia stages of Alzheimer’s disease^58^. Based on the involvement of P2YR, astrocytes were also identified as a target of interest to normalize network dysfunction in an Alzheimer’s disease mouse model^11, 43^. Indeed, at late disease stages, P2YR blockade was shown to regulate astrocytic and neuronal activity, preventing the decline in spatial memory observed in APP/PS1-21 mice^43^. Other strategies blocking the astrocytic adenosine A_2A_ receptor from an asymptomatic stage prevented spatial memory impairment and maintained low brain amyloid levels in the APPswe/PS1dE9 Alzheimer’s disease mouse model^59^. Compared to these encouraging strategies, one advantage of targeting TRPA1 is that this channel appears to be only mildly involved in maintaining physiological function. Thus, it seems to behave more as an “aggression sensor” in noxious conditions^15–17^. Indeed, the data presented here indicate that in healthy WT mice, chronic TRPA1 blockade had no effect on astrocyte activity, neuronal function, synapse structure or mnesic performance. Congruently, various TRPA1 antagonists have been successfully tested in human phase 1 studies and no adverse effects were reported^60, 61^. These results strengthen the relevance and attractiveness of TRPA1 as a therapeutic target.

The recurrent failure of therapeutic strategies for Alzheimer’s disease is partly due to the fact that they target late stages of the disease^21, 62^. The current boom in the search for diagnostic methods to identify early Alzheimer’s disease biomarkers^63^ or therapeutic targets involved in the prodromal phase (e.g., aMCI), must therefore be encouraged if we wish to prevent progression of neurodegeneration. The prodromal phase has been associated with hippocampal hyperactivity, thought to be an early, apparently reversible, neuronal dysfunction leading to future disease progression^57^. One of the major findings of our study is the primary role played by astrocytes in triggering this early neuronal hyperactivity. There is no doubt that this result represents a major conceptual shift in our understanding of how Alzheimer’s disease evolves, and that it should be taken into consideration when designing future therapeutic strategies. In conclusion, we propose that the TRPA1 channel could be an innovative target for the development of neuroprotective treatments.

## Acknowledgements

We gratefully acknowledge Magali Bartolomucci and the GIN behavioural facility, supported by the Grenoble Centre of Excellence in Neurodegeneration (GREEN). Our thanks to Dr Jean-Pierre Mothet for providing *R. gracilis* DAAO. We thank Dr Anthony Bosson, Dr Elodie Fino, Dr Muriel Jacquier-Sarlin and Dr Yves Goldberg for stimulating scientific discussion.

## Funding

This work was supported by INSERM, University Grenoble Alpes, France Alzheimer and the French National Research Agency in the framework of the “Investissements d’avenir” program (ANR-15-IDEX-02).

The laboratory hosting AP, SB, KPG, AB and MA is a member of the Grenoble Centre of Excellence in Neurodegeneration (GREEN).

## Competing interests

The authors report no competing interests.

## Supplementary material

Supplementary material is available at *Brain* online.

## Notes

### Competing Interest Statement

The authors have declared no competing interest.

