## Supplementary materials for "Astrocyte-neuron interplay is critical for Alzheimer’s disease pathogenesis and is rescued by TRPA1 channel blockade"

| Stage | Parameter | Condition | *p* value |
|---|---|---|---|
| **1-month-old** | sEPSCs frequency (Hz) | Tg male *vs* Tg female | 0.8072 |
| | sEPSCs amplitude (pA) | Tg male *vs* Tg female | > 0.999 |
| | Astrocyte active microdomains (%) | Tg male *vs* Tg female | 0.1481 |
| | Active microdomains frequency (calcium event /min) | Tg male *vs* Tg female | 0.2506 |
| **3-month-old** | sEPSCs frequency (Hz) | Tg male *vs* Tg female | 0.5934 |
| | sEPSCs amplitude (pA) | Tg male *vs* Tg female | 0.8851 |
| **6-month-old** | Time to escape box (s) | Tg male *vs* Tg female | 0.6029 |
| | Velocity (cm/s) | Tg male *vs* Tg female | 0.9998 |
| | Latency to target area (s) | Tg male *vs* Tg female | 0.8702 |

**Supplementary Table 1 Statistical analysis of the sex influence on studied parameters in APP/PS1-21 mice injected with vehicle** (Kruskal-Wallis test followed by Dunn's multiple comparisons test)

| Stage | Parameter | Condition | *p* value |
|---|---|---|---|
| **1-month-old** | sEPSCs frequency (Hz) | Tg male *vs* Tg female | 0.7108 |
| | sEPSCs amplitude (pA) | Tg male *vs* Tg female | 0.3841 |
| **3-month-old** | sEPSCs frequency (Hz) | Tg male *vs* Tg female | > 0.999 |
| | sEPSCs amplitude (pA) | Tg male *vs* Tg female | 0.3976 |
| | Astrocyte active microdomains (%) | Tg male *vs* Tg female | > 0.999 |
| | Active microdomains frequency (calcium event /min) | Tg male *vs* Tg female | 0.2358 |
| **6-month-old** | Time to escape box (s) | Tg male *vs* Tg female | 0.5585 |
| | Velocity (cm/s) | Tg male *vs* Tg female | 0.2062 |
| | Latency to target area (s) | Tg male *vs* Tg female | 0.2217 |

**Supplementary Table 2 Statistical analysis of the sex influence on studied parameters in APP/PS1-21 mice injected with HC030031** (Kruskal-Wallis test followed by Dunn's multiple comparisons test).

| **Stage** | **Parameter** | **Condition** | ***p* value** |
|---|---|---|---|
| **1-month-old** | sEPSCs frequency (Hz) | Tg *vs* Tg + vehicle | 0.7021 |
| | | WT vs WT + vehicle | 0.3437 |
| | sEPSCs amplitude (pA) | Tg *vs* Tg + vehicle | > 0.999 |
| | | WT vs WT + vehicle | > 0.999 |
| | Astrocyte active microdomains (%) | Tg *vs* Tg + vehicle | > 0.999 |
| | | WT vs WT + vehicle | > 0.999 |
| | Active microdomains frequency (calcium event /min) | Tg *vs* Tg + vehicle | 0.3879 |
| | | WT vs WT + vehicle | 0.9761 |
| **3-month-old** | sEPSCs frequency (Hz) | Tg *vs* Tg + vehicle | > 0.999 |
| | | WT vs WT + vehicle | 0.5131 |
| | sEPSCs amplitude (pA) | Tg *vs* Tg + vehicle | > 0.999 |
| | | WT vs WT + vehicle | 0.9271 |
| **6-month-old** | Time to escape box (s) | Tg *vs* Tg + vehicle | >0.999 |
| | | WT vs WT + vehicle | >0.999 |
| | Velocity (cm/s) | Tg *vs* Tg + vehicle | 0.2239 |
| | | WT vs WT + vehicle | 0.098 |
| | Latency to target area (s) | Tg *vs* Tg + vehicle | >0.999 |
| | | WT vs WT + vehicle | 0.2909 |

**Supplementary Table 3 Statistical analysis of the safety and inefficiency of the vehicle daily administration in APP/PS1-21 mice and their WT littermates** (Kruskal-Wallis test followed by Dunn's multiple comparisons test).

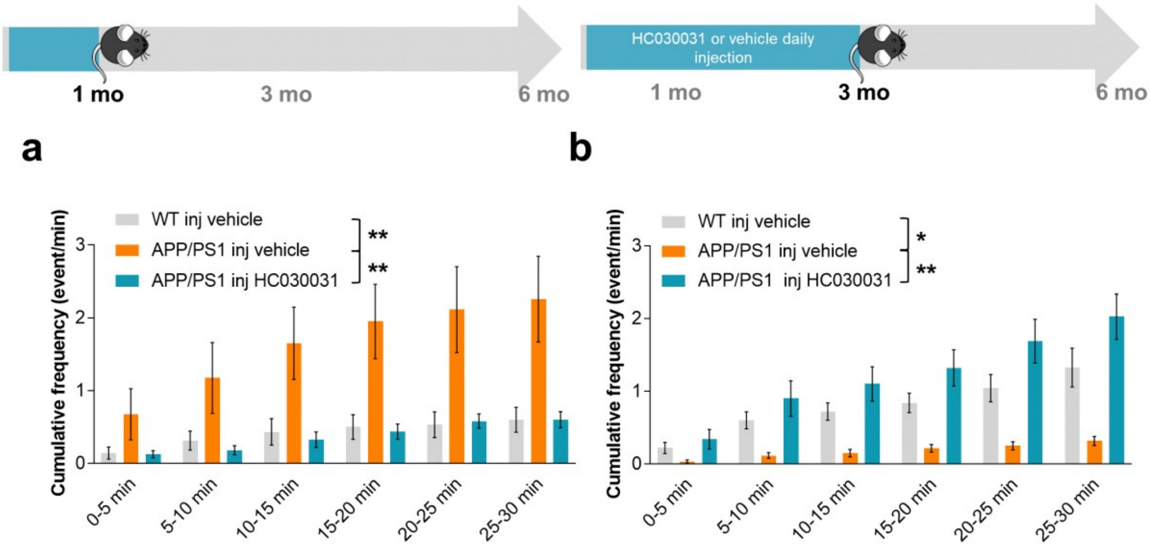


**Supplementary Figure 1 Chronic HC030031 treatment prevents additional astrocytic glutamate release in 1-month-old APP/PS1-21 mice and extrasynaptic activation of CA1 neurons. (A)** Time course of cumulative SIC frequency in vehicle-treated WT mice (grey; $n$ = 7 neurons from 4 mice), vehicle-treated APP/PS1-21 mice (orange; $n$ = 8 neurons from 5 mice), and HC030031-treated APP/PS1-21 mice (cyan; $n$ = 11 neurons from 5 mice) at 1-month-old. **(B)** Time course of cumulative SIC frequency in vehicle-treated WT mice (grey; $n$ = 10 neurons from 9 mice), vehicle-treated APP/PS1-21 mice (orange; $n$ = 12 neurons from 8 mice), and HC030031-treated APP/PS1-21 mice (cyan; $n$ = 12 neurons from 6 mice) at 3 months old highlighting a significant reduction in SIC frequency in vehicle-treated transgenic mice *vs* their WT littermates, in line with the observed reduction in the proportion of tripartite synapses. HC030031 treatment increased the SIC frequency to the same range as WT values. *, $p < 0.05$; **, $p < 0.01$; ***, $p < 0.001$ (Kolmogorov-Smirnov test).

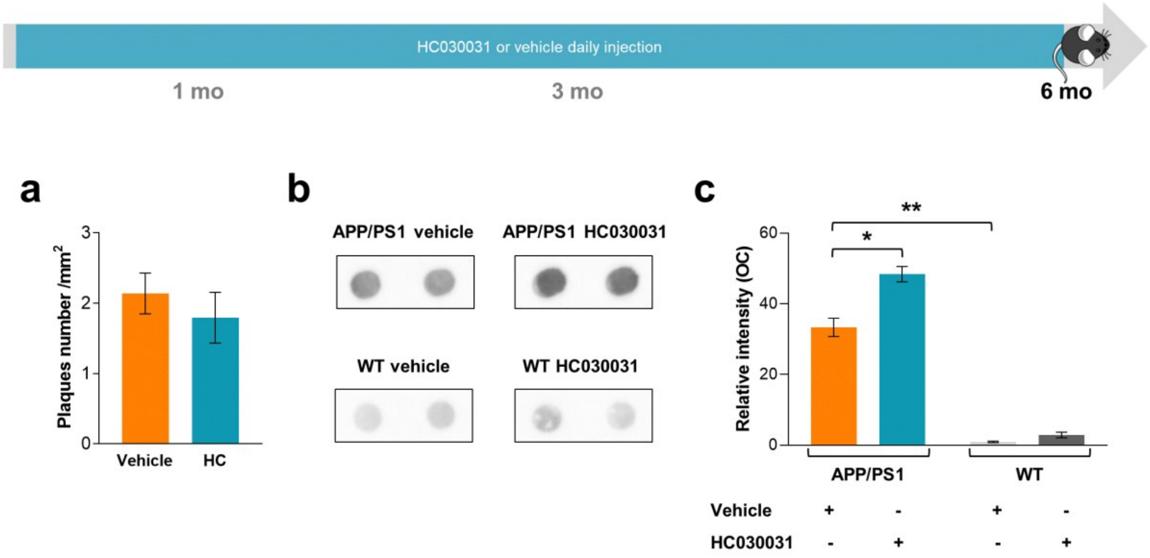


**Supplementary Figure 2 Chronic HC030031 treatment increases amyloid fibril compaction in 6-month-old APP/PS1-21 mice. (A)** Quantification of Thioflavin S-positive Aβ plaque numbers in the hippocampus from vehicle-treated (orange; $n = 18$ fields from 3 mice) and HC030031-treated (cyan; $n = 17$ fields from 3 mice) APP/PS1-21 mice. **(B)** Dot-blot assays with OC antibodies on hippocampal extracts from 6-month-old WT and APP/PS1-21 mice treated daily with either vehicle or HC030031 (2 extracts shown per condition). **(C)** Quantification of OC staining normalized relative to total protein loading shows an increase in the accumulation of Aβ fibrils in HC030031-treated transgenic mice ($n = 6$ hippocampi from 3 mice for each condition). *, $p < 0.05$; **, $p < 0.01$; ***, $p < 0.001$; n.s., not significant (Mann-Whiney or Kruskal-Wallis test).
